# Resolving haplotype variation and complex genetic architecture in the human immunoglobulin kappa chain locus in individuals of diverse ancestry

**DOI:** 10.1101/2023.10.23.563321

**Authors:** Eric Engelbrecht, Oscar L. Rodriguez, Kaitlyn Shields, Steven Schultze, David Tieri, Uddalok Jana, Gur Yaari, William Lees, Melissa L. Smith, Corey T. Watson

**Author notes:** Address correspondence to: Corey T. Watson, Melissa L. Smith.

## Abstract

Immunoglobulins (IGs), critical components of the human immune system, are composed of heavy and light protein chains encoded at three genomic loci. The IG Kappa (IGK) chain locus consists of two large, inverted segmental duplications. The complexity of IG loci has hindered effective use of standard high-throughput methods for characterizing genetic variation within these regions. To overcome these limitations, we leverage long-read sequencing to create haplotype-resolved IGK assemblies in an ancestrally diverse cohort (n=36), representing the first comprehensive description of IGK haplotype variation at population-scale. We identify extensive locus polymorphism, including novel single nucleotide variants (SNVs) and a common novel ∼24.7 Kbp structural variant harboring a functional IGKV gene. Among 47 functional IGKV genes, we identify 141 alleles, 64 (45.4%) of which were not previously curated. We report inter-population differences in allele frequencies for 14 of the IGKV genes, including alleles unique to specific populations within this dataset. Finally, we identify haplotypes carrying signatures of gene conversion that associate with enrichment of SNVs in the IGK distal region. These data provide a critical resource of curated genomic reference information from diverse ancestries, laying a foundation for advancing our understanding of population-level genetic variation in the IGK locus.

## Introduction

Immunoglobulins (IGs) are critical protein components of the immune system with roles in both innate and adaptive responses [1]. IGs are produced by B lymphocytes and are either expressed on the cell membrane as B cell receptors (BCRs) or secreted as antibodies. IGs are composed of two pairs of identical ‘heavy’ chains and ‘light’ kappa or lambda chains, encoded by genes located at three loci in the human genome: the IG heavy chain locus (IGH) at 14q32.33, and the IG lambda (IGL) and kappa (IGK) loci, located at 22q11.2 and 2p11.2, respectively [2]. Through the unique mechanism of V(D)J recombination [3], individual variable (V), diversity (D) and joining (J) genes at the IGH locus, and V and J genes at either the IGK or IGL loci, somatically rearrange to generate V-D-J and V-J regions that encode respective antibody variable domains. IG loci are enriched for large structural variants (SVs), including insertions, deletions, and duplications of functional genes, many of which are variable between human populations [4–7]. The IGH and (to a lesser extent) IGL loci exhibit extensive haplotype diversity and structural complexity, which contributes to challenges associated with IG haplotype characterization using standard high-throughput approaches [4,6–9]. This has limited our understanding of the impact of IG genetic variation in immune function and disease [10–12].

A unique structural characteristic of the IGK locus is that it includes two large inverted segmental duplications (SDs) that contain distinct V genes; these paralogous regions (termed proximal and distal) [13,14] are separated by an array of 45-Kbp highly identical repeats that encode the 45*S* rRNA [15]. The proximal region spans 500 Kbp and includes 69 IGKV functional/ORF genes and pseudogenes, downstream of which are 5 functional IGKJ genes and a single functional IGK constant gene. The distal region spans 430 Kbp and includes 62 V genes and pseudogenes (distal V genes are denoted by a ‘D’; for example, IGKV1D-12) [2]. An additional unmapped IGKV gene has also been reported [16] and is cataloged in the International Immunogenetics Information System (IMGT) [17] as IGKV1-NL1 (where the NL indicates “not localized”). The documentation of this gene suggests the presence of SVs in IGK, consistent with observations in IGH and IGL [4,6,7,14]. Evidence for sequence conversion between proximal and distal regions has also been reported [14], highlighting the potential for complex genetic signatures to be observed at the population level.

To date, there are 108 functional/ORF IGKV, IGKJ, and IGKC alleles curated in IMGT. However, only a limited number (n=4) of fully curated and annotated haplotypes have been characterized at the genomic level [13,14,18], and adaptive immune receptor repertoire sequencing (AIRR-seq) has indicated that allelic variation in IGK is likely to be more extensive than what is currently documented [19,20]. Compared to IGH [4,6] and IGL [7], our understanding of germline variation at the IGK locus is limited, especially in terms of haplotype structure (including structural variation) as well as coding and non-coding polymorphism. It will be critical to further catalog this genetic variation if we are to clarify the impact of IG germline variants on the antibody response. Diversity in antigen-naïve B cell antibody repertoires is due to combinatorial and junctional diversity and the effective pairing of particular heavy and light chain genes [1]. This partly ensures that the immune system is able to recognize and mount effective immune responses against a broad range of potential antigens. Critically, however, population-level IG haplotype and allelic variation also makes important contributions to antibody diversity and function [6,11,21,22], including in the context of infection and vaccine responsiveness. For example, there have now been clear demonstrations that IGH locus SVs and non-coding single nucleotide variants (SNVs) impact usage of IGH genes in expressed antibody repertoires [6]. Antibody light chains also contribute to antigen binding and can limit self-reactivity through the process of light chain receptor editing during B cell development (reviewed in [23]). Consistent with findings related to IGH diversity, usage of specific IGKV genes has been associated with neutralizing antibodies against protein components of influenza A (IGKV4-1, IGKV3-11, IGKV3-15; [24,25]), HIV-1 (IGKV3-20, IGKV1-33; [26]), Zika virus (IGKV1-5; [27]), as well as auto-reactivity in celiac disease (IGKV1-5; [28]) and systemic lupus erythematosus (IGKV4-1; [29]). Taken together, these findings further motivate a need to develop more robust methods for detecting and genotyping IGK polymorphisms.

To characterize germline IGK sequence variation at population-scale, we extended a technique that employs targeted long-read sequencing to generate haplotype-resolved assemblies of IGK proximal and distal regions [6–8]. Here, we characterize IGKC, IGKJ, and IGKV alleles in 36 individuals from the 1000 genomes project (1KGP) [30]. We report 64 novel IGKV alleles and provide evidence of inter-population variation in IGKV allele frequencies. We identify a structural variant (SV) that includes a previously unlocalized IGKV gene (IGKV1-NL1) and provide evidence that this SV is common across populations. In addition, we report that a previously described gene conversion event [14] is associated with haplotype structure in the IGK distal region.

## Material and Methods

### Sample information

Samples used in this study were previously collected as part of the 1KGP [30]. Genomic DNA isolated from lymphoblastoid cell lines (LCLs) were procured for all individuals from Coriell Repositories (https://www.coriell.org/; Camden, NJ). Sample population, subpopulation, and sex are reported in **Table S1**.

### Construction of a custom reference assembly for the IGK locus

The GRCh38 assembly gap between the IGK proximal and distal regions was modified as follows: 10 Kbp of sequence in the GRCh38 assembly upstream of IGKV2-40 and downstream of IGKV2D-40 was swapped with homologous sequence in the T2T assembly [15]; the remaining 190,249 bases in the GRCh38 assembly gap were N-masked. 24,729 bp in a hifiasm-generated assembly for the sample HG02433 were inserted between IGKV1D-8 and IGKV3D-7; 10 Kbp of sequence flanking the insertion in this assembly were swapped with homologous sequence in GRCh38.

Coordinates of IGK gene features (L-Part1, Intron, V-exon, RSS) were determined using the program ‘Digger’ (https://williamdlees.github.io/digger/_build/html/index.html) and will be made available, along with the custom reference, at https://vdjbase.org/.

### IGK sequencing and phased assembly

To enrich for DNA from the IGK locus, we designed a custom Roche HyperCap DNA probe panel (Roche) that includes target sequences from the GRCh38 IGK proximal (chr2:88,758,000-89336429) and distal (chr2:89846189-90,360,400) regions. Genomic DNA samples were prepared and sequenced following previously described methods [16]. Briefly, genomic DNA was sheared to ∼16 Kbp using g-tubes (Covaris, Woburn, MA, USA) and size-selected using the Blue Pippin system (Sage Science, Beverly, MA, USA), collecting fragments that ranged in size between 11.9 – 20.6 Kbp (mean=15.2 Kbp). Size-selected DNA was ligated to universal barcoded adapters and amplified, then individual samples were pooled in groups of six prior to IGK enrichment using the custom Roche HyperCap DNA probes [16]. Targeted fragments were amplified after capture to increase total mass, followed by SMRTBell Express Template Prep (Kit 2.0, Pacific Biosciences) and SMRTBell Enzyme Cleanup (Kit 2.0, Pacific Biosciences), according to the manufacturer’s protocol. Resulting SMRTbell libraries were pooled in 12-plexes and sequenced using one SMRT cell 8 M on the Sequel IIe system (Pacific Biosciences) using 2.0 chemistry and 30 hour movies. Following data generation, circular consensus sequencing analyses were performed, generating high fidelity (“HiFi”, >Q20 or 99.9%) reads for downstream analyses. Reads aligned to the custom GRCh38 reference were input to bamtocov [31] for coverage analysis (**Figure S1**).

HiFi reads were processed using IGenotyper [8]; the programs “phase” and “assemble” were run with default parameters to generate phased contigs and Hifi read alignments to the custom IGK reference assembly. HiFi reads were also used to generate haplotype-phased *de novo* (i.e. reference-agnostic) assemblies using hifiasm [32] with default parameters. Hifiasm assembly contigs were mapped to the custom IGK reference using minimap2 (v2.26) with the ‘-x asm20’ option. For each sample, aligned HiFi reads as well as aligned IGenotyper-generated contigs and hifiasm-generated contigs were viewed in the Integrative Genomics Viewer (IGV) application for manual selection of phased contigs. Where a hifiasm and an IGenotyper contig were identical throughout a phased block, the hifiasm contig was selected (example: **Figure S4**). Examples of contig selection are provided in **Figures S4-S7**. Curated, phased assemblies were aligned to the custom IGK reference using minimap2 with the ‘-x asm20’ option. Phased assembly coverage across IGK (**Figure S2**) was computed using ‘bedtools coverage’ (bedtools v2.30.0) [33]. Average IGenotyper and hifiasm contig lengths across IGK (**Figure S1**) were computed by extracting contig lengths using ‘samtools view’.

To assess accuracy of assemblies, IGK-personalized references were first generated by N-masking the IGK locus (chr2:88837161-90280099) and, for each sample, appending the reference FASTA file with IGK curated assembly sequences. Hifi reads from each individual were aligned to the corresponding IGK- personalized reference using minimap2 with the ‘-x map-hifi’ preset. Positions in IGK assembly contigs with > 25% of aligned HiFi read mismatch were identified by parsing the output of samtools ‘mpileup’ using a custom script (https://github.com/Watson-IG/swrm_scripts/blob/master/ee/assembly_analysis_IGK/perbase_accuracy.sh); assembly accuracy was determined using the formula [total mismatch bases / total (diploid) assembly length (bp)] * 100 = % accuracy (**Table S2**).

### Genotyping and analysis of IGK assemblies

Sample IDs were inserted into the @RG tag of curated assembly BAM files using the ‘samtools addreplacerg’ command (samtools v1.9) [25]. Variants were called from assemblies using the ‘bcftools mpileup’ (bcftools v1.15.1) command with options ‘-f -B -a QS’ and a ‘--regions’ (BED) file corresponding to the IGK locus (proximal: chr2:88866214-89331428, distal: chr2:89851190-90265885), then the ‘bcftools call’ command with the ‘-m’ option. Multiallelic SNVs were split into biallelic SNVs using the ‘bcftools norm’ command with options ‘-a -m-’. A mutli-sample VCF file was generated using ‘bcftools merge’ with the ‘-m both’ option, and the INFO field annotations for V-exon, introns, L-Part1, RSS, and intergenic sequences were added using vcfanno [34]. The VCF file was filtered to include SNVs with an alternate allele count > 1 (MAF ≥ 2.86%) using ‘bcftools view’ with the ‘-v snps -i ’INFO/AC > 1.0’ options’.

The filtered VCF file was input for PCA of SNVs in the IGK distal region (chr2:89852177-90266726) using the SNPRelate package [35] command ‘snpgdsVCF2GDS’ with the option ‘method = “biallelic.only”’ followed by the ‘snpgdsPCA’ command with default parameters. The SNPRelate command ‘snpgdsHCluster’ was used for hierarchical clustering of samples using distal region SNVs as input, for visualization of genotypes in **Figure 3E**. A parsable genotype table (**Table S9**) was generated from the filtered VCF using ‘bcftools query’ and read into R for analysis (**Figure 2A, 2C-D**, **Figure 3D-F**). SNV densities over genomic intervals were calculated using the formula: SNV density = [sum of alternate alleles over interval / interval length (bp) * 2]. 10 Kbp intervals along the IGK distal region were generated using the ‘bedtools makewindows’ [33] command (**Table S10**, **Table S11**). dbSNP data were downloaded from the UCSC Table Browser [36] (dbSNP ‘common variants’ release 153). The 1KGP “phase 3” GRCh38 chromosome 2 multi-sample VCF file (https://ftp.1000genomes.ebi.ac.uk/vol1/ftp/data_collections/1000G_2504_high_coverage/working/phase3_lifto <u>ver_nygc_dir/phase3.chr2.GRCh38.GT.crossmap.vcf.gz</u>) was filtered to include only the IGK locus (chr2:88866214-90265885) excluding the intervening gap region (chr2:89331429-89851189), split into single- sample files, filtered to include only SNVs using bcftools ‘view’, and compared with SNVs from curated assemblies using bedtools ‘intersect’. The downloaded multi-sample VCF file included 33 of 35 samples used in this study for analyses presented in **Figures 2-4**. Both dbSNP and 1KGP coordinates were lifted over to our custom reference to account for the SV insertion.

**Figure 1.**
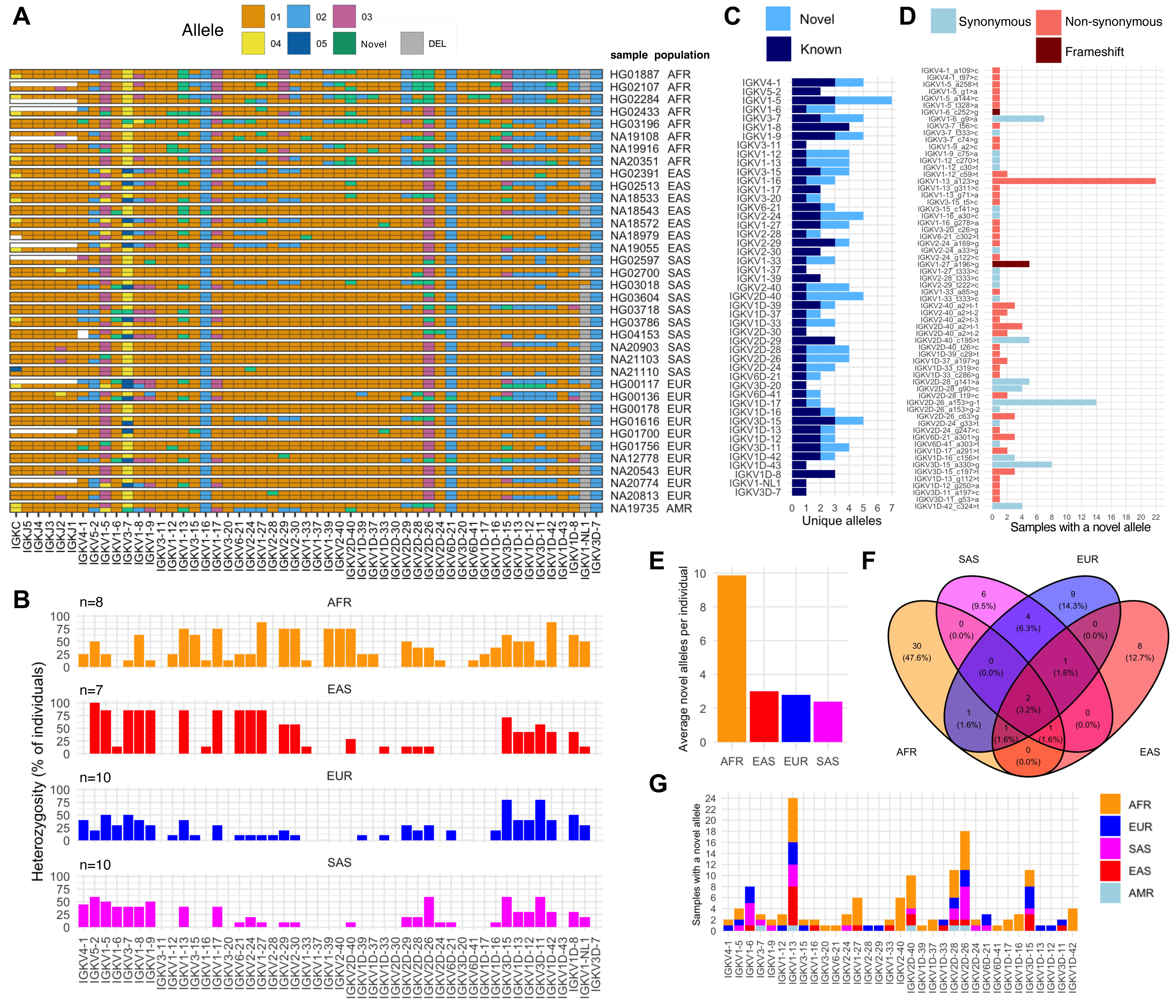
Characterization of known and novel IGK alleles in a cohort of 36 individuals. **(A)** Diagram of alleles for the single IGKC gene, 5 IGKJ genes, and 47 IGKV genes. Individuals (rows) are split into two sub-rows, each indicating the allele for a gene (column). Tile colors correspond to allele sequences cataloged in the IMGT database, which have numerical designations. Allele sequences absent from IMGT are considered novel and colored lime-green. Gray tiles represent allele absence due to structural variation (deletion); white tiles represent regions unresolveable due to V(D)J recombination artifacts, which we have shown previously to occur in DNA derived from some LCLs [7,8]. Phased assemblies were created for the proximal and distal regions separately; individual sub-rows do not represent locus-wide haplotypes. **(B)** Bar plot of the percent of individuals in each population that are heterozygous for 47 IGKV genes. **(C)** Stacked bar plot of the number of known and novel alleles for each of 47 IGKV genes; at least one novel allele was identified for 33 of these genes. **(D)** Bar plot of the number of individuals that have at least one allele for each of 64 novel alleles. Each novel allele is colored to indicate the presence of a) only synonymous, b) at least one non-synonymous, or c) a frame-shift substitution. **(E)** Bar plot of the average number of novel alleles identified in each population. **(F)** Venn diagram illustrating the distribution of 64 novel alleles among populations. **(G)** Stacked bar plot of the number of individuals (colored by population) with at least one novel allele for each of 33 IGKV genes.

**Figure 2.**
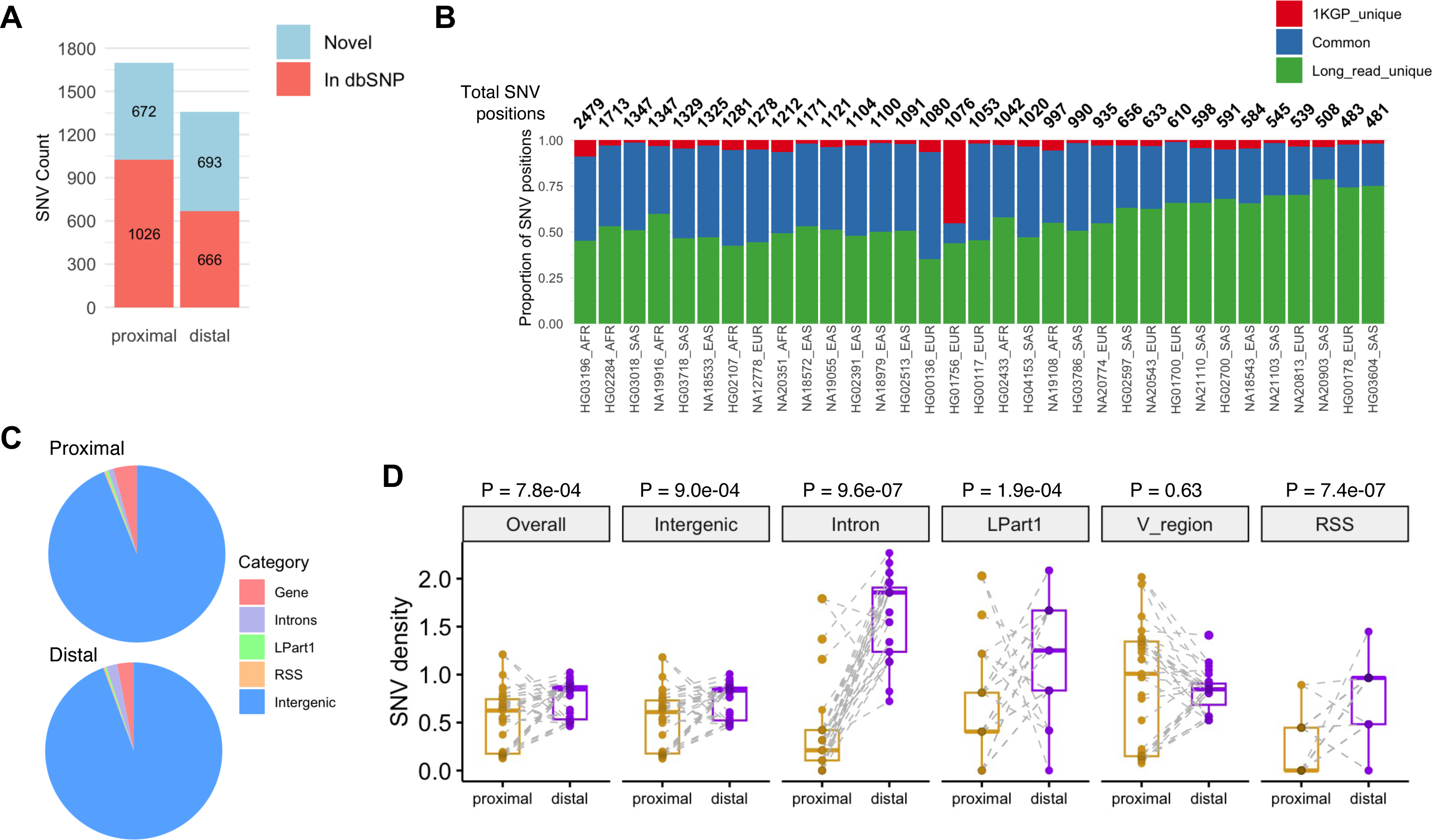
Single nucleotide variation in IGK proximal and distal regions. **(A)** Stacked bar plot of the number of SNVs (MAF ≥ 2.86%) in the IGK proximal and distal regions. Bar colors indicate SNV inclusion in dbSNP. **(B)** Stacked bar plot of IGK locus SNV positions found uniquely in the 1KGP “phase 3” release, uniquely in curated assemblies generated in this study (long read-based assemblies), or in both, for 33 individuals. Total SNV positions are indicated above bars. **(C)** Pie charts of SNV distribution among genic (functional IGKV genes) and intergenic regions in proximal and distal regions of IGK. **(D)** Boxplots of SNV densities in proximal and distal regions for each individual (data point). Gray dashed lines indicate paired SNV density values for individuals. Gray shading of data points indicates data point overlap. P- values: paired Wilcoxon tests.

**Figure 3.**
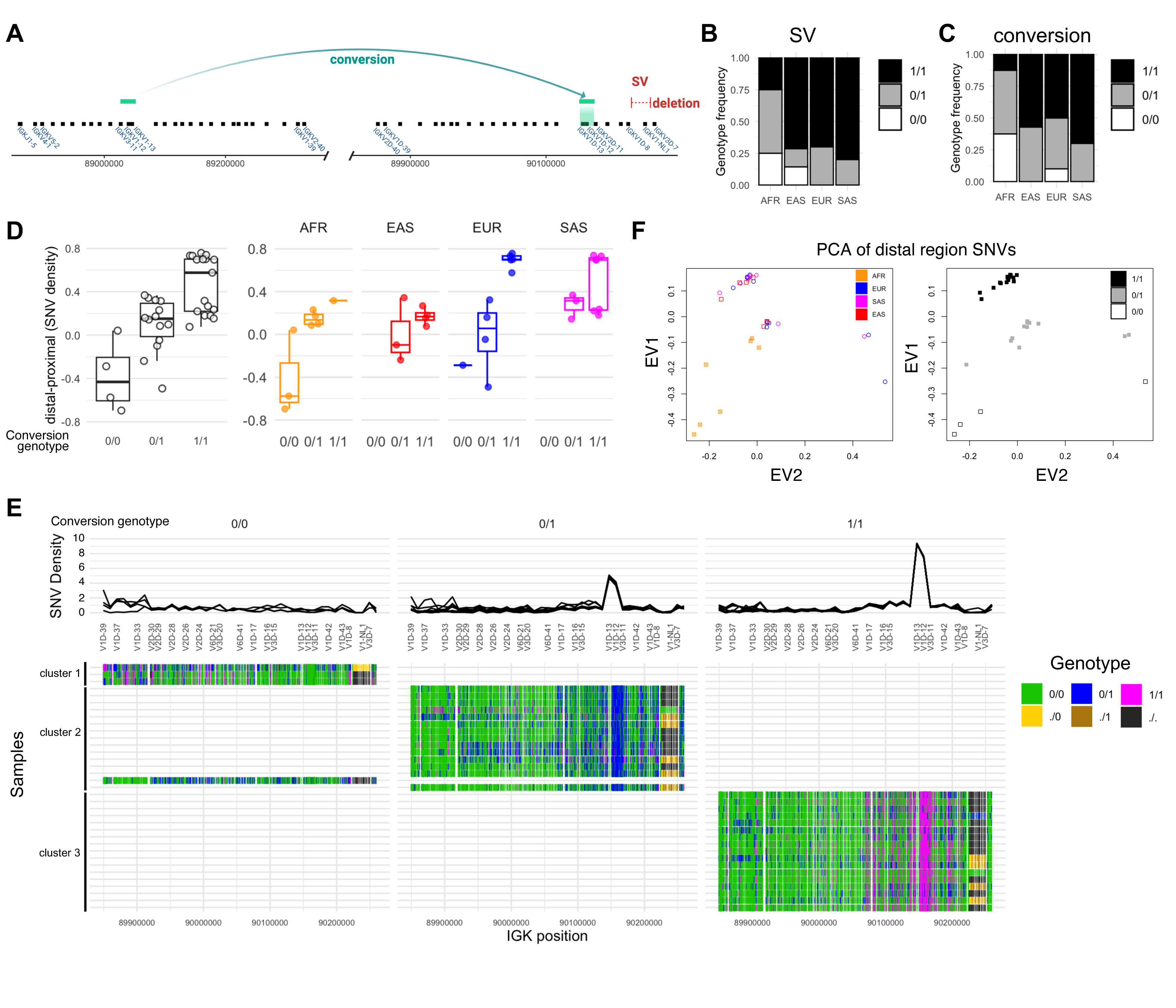
A gene conversion event and structural variation are common in the IGKV distal region **(A)** Diagram of a gene conversion event in which ∼16 Kbp of sequence containing IGKV1-12 and IGKV1-13 replaced paralogous sequence in the distal region, described previously [14]. A 24.7 Kbp deletion SV that includes the gene IGKV1-NL1 is also shown. **(B)** Stacked bar plot indicating the SV genotype frequencies for each population, with “1” corresponding to deletion. **(C)** Stacked bar plot indicating the gene conversion haplotype frequency for each population, with “1” corresponding to the conversion haplotype. **(D)** Boxplot of the SNV density difference between distal and proximal regions with samples grouped according to their genotype for the gene conversion (left) as well as population (right). **(E)** (Top) SNV densities in 10 Kbp windows along the IGK distal region (chr2:89859172-90266726). IGKV genes in this interval are labeled along the x-axis. Each line represents a sample. Samples are grouped according to their genotype for the gene conversion. (Bottom) Genotypes (colors) at each variant position for the corresponding coordinates above. Hierarchical clustering of samples (rows) resulted in the 3 indicated clusters. **(F)** Principal component analysis of samples based on SNVs (MAF ≥ 2.86%) in the IGK distal region. Samples are colored by population (left) or gene conversion genotype (right). EV1: eigenvector 1, EV2: eigenvector 2

**Figure 4.**
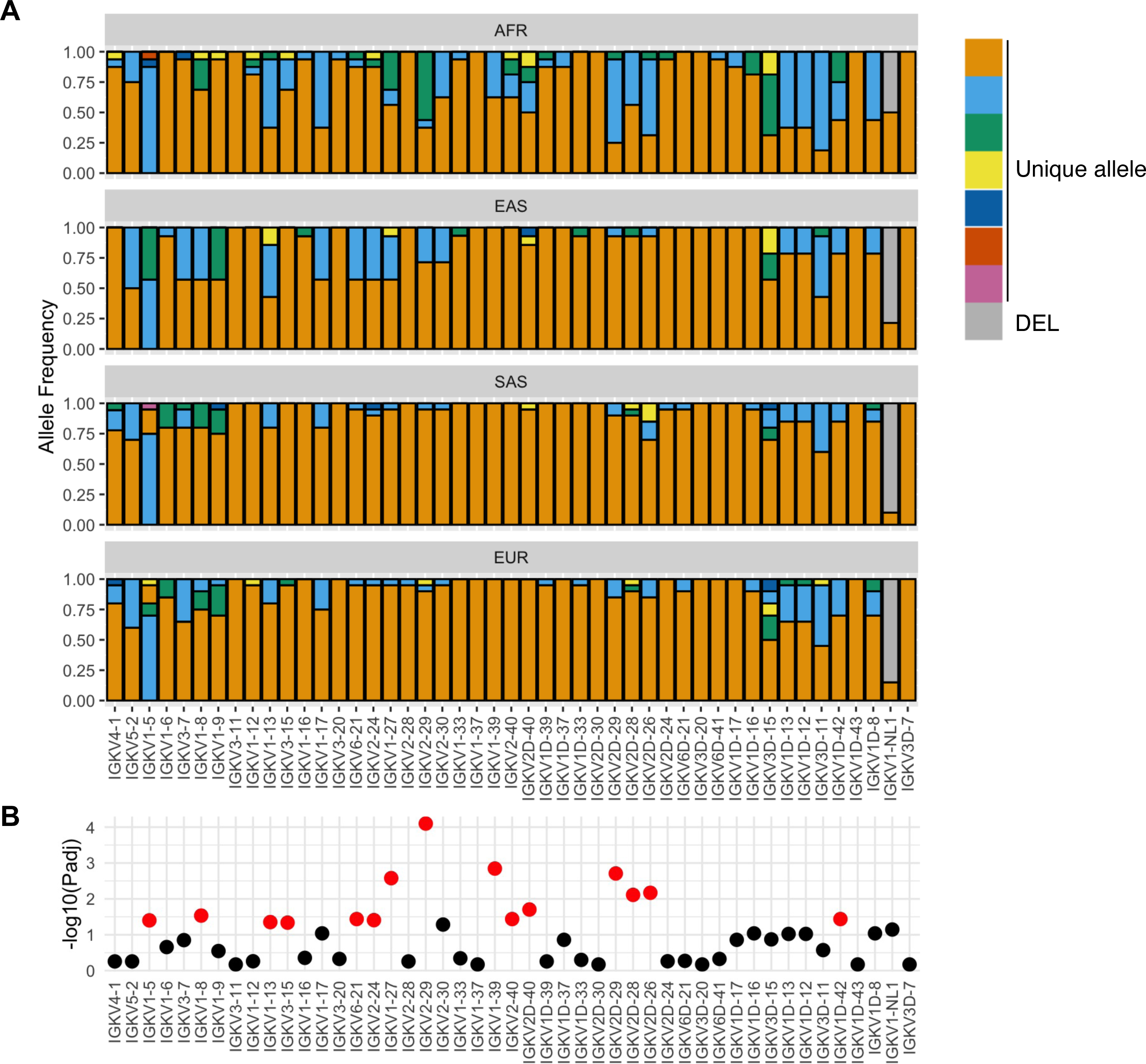
Inter-population IGKV allelic variation **(A)** Stacked bar plot of allele frequencies for each of 47 functional IGKV genes for AFR, EAS, SAS, and EUR populations. For each gene, a color corresponds to a unique allele (detailed in **Table S5**). Gray color indicates allele absence due to structural variation (deletion). **(B)** Differences in allele frequency distributions among populations were determined using a chi-square test (**Table S6**); Benjamini-Hochberg adjusted P-values (Padj) are plotted as -log10(Padj) for each gene. Red points indicate Padj < 0.05.

### Identification of IGKC, IGKJ, and IGKV alleles

Allele sequences at V-exons were extracted from manually curated phased assemblies using a custom python script (https://github.com/Watson-IG/swrm_scripts/blob/master/ee/assembly_analysis_IGK/extract_sequence_from_bam_EEmod3.py) to generate a FASTA file of alleles for each sample. Per-sample allele FASTA files were queried against all human IGK alleles cataloged in IMGT [17] (retrieved 2022/11/21) using BLAT [37] as follows: ‘blat <IMGT_alleles.fasta> <sample_alleles.fasta> -out=blast8’. Alleles from samples were assigned to the IMGT- cataloged allele with the lowest e-value in the BLAT output; sample allele sequences with < 100% identity and at least 1 mismatch relative to the best-matching (lowest e-value) IMGT allele were defined as novel (**Table S3**). Novel alleles were analyzed using IMGT/V-QUEST [38] to determine the presence of synonymous, non-synonymous, and frame-shift substitutions.

Novel alleles were examined for HiFi read support by alignment of HiFi reads to an IGK-personalized reference for each sample. **Figure S10** includes an example; IGV screenshots of HiFi reads and novel alleles from the sample HG01887 aligned to the corresponding IGK-personalized reference using minimap2 with the ‘- x map-hifi’ option.

### Statistical Analyses

Statistical tests described in Results and Figure Legends associated with **Figure 2** (**Table S9**), **Figure S8** (**Table S9**), **Figure 3** (**Table S6**), and **Figure 4** (**Table S7**, **Table S8**), were performed in R (v4.2.1).

## Results

### Assembly of the IG kappa chain locus

To sequence the IGK locus, we extended our previously described method [6–8] such that genomic DNA fragments (∼15 Kbp in length) derived from this locus were enriched using targeted probes and sequenced using single-molecule, real-time (SMRT) long-read sequencing. This methodology was applied to an ancestrally diverse cohort (n=36; **Table S1**) to assess genetic variants in IGK across different populations. The cohort represented individuals of African (AFR, n=8), East Asian (EAS, n=7), South Asian (SAS, n=10), European (EUR, n=10), and Central American (AMR, n=1) ancestry, spanning 18 subpopulations. Average high quality (>99% accuracy, “HiFi”) read coverage across the custom IGK reference (see **Materials and Methods**) was > 50X within the proximal and distal IGK regions for all samples (**Figure S1**). HiFi reads were assembled into phased contiguous sequences (contigs) using both IGenotyper [6,8], which uses reference-aligned reads to generate contigs, and hifiasm [32], which generates a *de novo* diploid assembly (**Figure S1**). The resulting phased contigs were assessed for read support and merged (see **Materials and Methods**) into manually-curated diploid assemblies (see **Materials and Methods** and **Figures S4-S7**), which on average spanned 99.3% (479.53 Kbp) and 99.7% (413.26 Kbp) of the proximal and distal regions, respectively (**Figure S2**). Accuracies of manually-curated diploid assemblies were determined by alignment of HiFi reads to IGK- personalized references (see **Materials and Methods**) and ranged from 99.63% to 99.97% (mean: 99.95%) (**Table S2**). Assembly contigs could not be phased between proximal and distal regions due to the extended non-IGK sequence region not covered by our probe panel; additionally, we cannot fully account for the possibility of rare phase-switch errors present in hifiasm assemblies. However, all assemblies were locally phased over each variant position, allowing us to effectively genotype IGK gene loci and locus-wide polymorphisms. For IGK gene and allele annotation, as well as variant calling, curated assemblies were aligned to our in-house reference assembly, which included a novel SV insertion identified in this cohort (see also **Materials and Methods**).

### Characterization of IGKV alleles

Allele sequences were annotated and extracted from haplotype-resolved sequences of all donors for 47 functional/ORF IGKV genes. These alleles were compared against those cataloged in IMGT [17], revealing both known (n=79) and novel (n=64) alleles (**Figure 1A, Tables S3-4**). Of the 64 novel alleles, 21 were observed in >1 individual; 9 of the 64 alleles were also independently supported by alleles curated in VDJbase [39] (**Table S3**). Genes with a single identified allele were determined to be homozygous across all individuals; these genes included IGKV3-11, IGKV1-37, IGKV2D-30, IGKV3D-20, IGKV1D-43, and IGKV3D-7 (**Figures 1A, 1B**). IGKV genes that were heterozygous in at least half of all individuals included IGKV5-2, IGKV1-5, IGKV1-8, IGKV1-13, IGKV1-17, IGKV1-27, IGKV3D-15 and IGKV3D-11 (**Figure 1B**). Heterozygosity varied among populations for several genes, including IGKV3-15, IGKV1-27, IGKV2-29, IGKV2-30, IGKV1-39, IGKV2-40, and IGKV2D-40 (**Figure 1B**, further discussed in **Figure 4**).

Novel alleles were identified for 17 of 24 proximal genes and 16 of 23 distal genes (**Figure 1C**) (**Table S3**). Among the 21 novel alleles identified in >1 individual, 8, 12, and 1 resulted in synonymous, non-synonymous, and frame-shift substitutions, respectively (**Figure 1D**). Thirty-three individuals (91.7%) carried at least one novel allele, with individuals of African descent harboring ∼10 novel alleles per individual on average (range: 6-13) and individuals from other populations harboring ∼2-3 novel alleles on average (ranges; EAS: 1- 5, SAS: 0-6, EUR: 0-6) (**Figure 1E**). Fifty-three (83%) of the novel alleles were exclusive to a single population, of which, 30 were identified only in AFR individuals (**Figure 1F**). Ten of the novel alleles were identified in two or more populations (**Figure 1F**).

Among all functional/ORF IGKV genes, the number of individuals harboring at least one novel allele was highest for IGKV1-13 (22 individuals), IGKV2D-26 (18 individuals), IGKV2D-28 and IGKV3D-15 (11 individuals each), followed by IGKV2D-40, IGKV1-6, IGKV2-40, IGKV1-27, IGKV1D-42, and IGKV1-5 (> 3 individuals each) (**Figure 1G**).

### Single nucleotide polymorphism in the IGKV proximal and distal loci

In addition to variants within IGKV V-exon regions, we also characterized SNVs across the whole of the IGK locus. We enumerated 3,057 SNVs for which we observed >1 individual carrying an alternate allele relative to our reference (equivalent to a MAF ≥ 2.86%). Of these, 40% and 51% were absent from the dbSNP ‘common variants’ dataset in the proximal and distal regions, respectively (**Figure 2A**). Sample-level comparison of SNV positions identified in this study with those from short read-derived (1KGP) data revealed that, on average, 55.8% of SNV positions were uniquely identified in our curated assemblies and 39.5% were identified in both datasets (**Figure 2B, Table S5**), consistent with known limitations of short reads for genotyping genomic regions enriched with duplicated sequence [8,10,40–42]. SNV positions were more numerous in individuals of non-EUR ancestry, most notably among individuals of AFR ancestry (**Figure 2B**).

We categorized the 3,057 SNVs as overlapping either intergenic sequence or an IGKV gene feature, including the leader part 1 (L-Part1), intron, V-exon, or recombination signal sequence (RSS). While the majority of SNVs were intergenic (**Figure 2C**), SNV density was greater in V-exons, L-Part1 regions, and introns relative to intergenic and overall SNV densities (**Figure S8**). Partitioning of SNVs into proximal and distal also revealed that, for each feature except V-exons, there was increased SNV density in the distal relative to proximal region for the majority of (but not all) individuals (**Figure 2D**).

### Characterization of a common structural variant, gene conversion haplotype and associated SNV signatures in the IGKV distal region

We analyzed our curated assemblies for evidence of large structural events. This revealed the presence of a ∼24.7 Kbp insertion between IGKV1D-8 and IGKV3D-7 in a large fraction of haplotypes in our cohort (**Figure S3**); this insertion is not present in GRCh38. The insertion sequence was queried against alleles from the IMGT database using BLAT [37], revealing the presence of a single functional IGKV gene, IGKV1-NL1 (IGKV1-“not localized 1”); this sequence was integrated into the reference assembly used for all analyses in this study (see **Materials and Methods**) and its absence was codified as a deletion (**Figure 3A, Table S1**). The frequency of the deletion was 50% in AFR as compared to 78.6%, 90%, and 85% in EAS, SAS, and EUR populations, respectively (**Figure 3B**). The distribution of this SV allele in AFR versus non-AFR groups was significantly different (Fisher’s exact test, *P* < 0.01) (**Table S6**). Among 35 individuals, one EAS and two AFR samples were identified as homozygous for the IGKV1-NL1 allele (**Figure 1A**).

Watson et al. (2015) [14] previously reported a gene conversion event wherein a ∼16 Kbp region that includes IGKV1-13 was swapped into the paralogous distal region (**Figure 3A**). Relative to a reference sequence that does not contain this sequence exchange, a haplotype with the gene conversion is marked by high SNV density within a ∼16 Kbp window that spans IGKV1D-12 and IGKV1D-13 (**Figure S6**). Genotyping of this gene conversion event revealed that it is a common haplotype structure in the population, with a frequency ≥ 70% in EUR, SAS, and EAS populations, and a frequency of 37.5% in AFR populations (**Figure 3C, Table S1**). The distribution of the gene conversion haplotype in AFR versus non-AFR groups was significantly different (Fisher’s exact test, *P* < 0.01) (**Table S6**). In this cohort, the four individuals homozygous for the reference (non-conversion) haplotype (HG01887, HG02107, HG02284, HG00136) were also the only four individuals homozygous for IGKV1D-13*02, IGKV1D-12*02, and IGKV1D-11*02 (**Figure 1A**), which may indicate that the gene conversion haplotype structure extends beyond the previously identified ∼16 Kbp window.

Based on the observation that most, but not all, individuals exhibit increased SNV density in the distal relative to proximal region (**Figure 2C**), we computed the within-individual difference between distal and proximal SNV densities and then partitioned individuals by genotype for the gene conversion event (**Figure 3D**). This analysis revealed that the gene conversion haplotype was associated with increased SNV density in the distal region (**Figure 3D**). Accordingly, SNV density in the region spanning IGKV1D-12 and IGKV1D-13 increases with dosage of the gene conversion haplotype (**Figure 3E**). We noted increased SNV density throughout an ∼70 Kbp region downstream of IGKV1D-13 extending to IGKV1D-17, and also throughout an ∼30 Kbp region from IGKV1D-11 to downstream of IGKV1D-43 (**Figure 3E**), further suggesting that the gene conversion haplotype structure is extended beyond the ∼16 Kbp window containing IGKV1D-12 and IGKV1D-13. Principal component analysis (PCA) based on these distal region SNVs revealed separation of individuals based on their genotype for the gene conversion haplotype (**Figure 3E**). Taken together, these data indicate that the gene conversion haplotype is a major driver of genetic variation in the IGK distal region.

### Inter-population IGKV allelic variation

Coding sequences in the human IGH and IGL loci have been shown to exhibit population-level allelic diversity [6,7,11,23]. Although our cohort is relatively small in number, we observed that functional/ORF IGKV genes also exhibited inter-population differences in allele frequency (**Figure 4A, Table S7**). In total, 15 of the 47 IGKV genes demonstrated significant skewing in allele frequencies (**Figure 4B**; chi-square, *P* < 0.05). For example, the alleles IGKV1-5*04 and IGKV1-8*02 were not identified in AFR or SAS individuals, but each had an allele frequency of 0.43 and 0.1 in EAS and EUR individuals, respectively (**Figure 4A**). The alleles IGKV6- 21*02 and IGKV2-24*02 were similarly enriched in EAS individuals (**Figure 4A**). These inter-population differences in allele frequency appeared to correlate with overall SNV density as six of seven EAS individuals showed SNV density patterns between IGKV6-21 and IGKV1-27 that were highly similar and, on average, distinct from AFR, EUR, and SAS populations (**Figure S9**). Alleles identified only in AFR individuals included IGKV3-15*02, IGKV1-39*02, three novel IGKV2-40 alleles, a novel IGKV2D-28 allele (IGKV2D-28_N_g141>a), and a novel IGKV1D-42 allele (IGKV1D-42_N_c324>t) (**Figure 4A**). We noted that SNV density was uniquely elevated for half (4/8) of AFR individuals in a region including IGKV1-39 and IGKV2-40 (**Figure S9**), suggesting a relationship between haplotype structure and gene alleles in this region. The frequencies of the novel alleles IGKV2D-26_N_a153>g and IGKV2D-29*02 were > 4-fold higher in AFR relative to each other population (**Figure 4A-B**). In contrast, the frequency of IGKV2D-29*01 was 0.25 in AFR and ∼0.9 in EUR, SAS, and EAS populations. The novel allele IGKV1-13_N_a123>g was most frequent in AFR individuals and the only two homozygous individuals (HG01887 and HG02107, **Table S3**) were both AFR and were also homozygous for the non-conversion (reference) haplotype (**Table S1**). The frequency of each previously highlighted allele exhibited significant inter-population variance as determined by chi-square test (**Figure 4B, Table S8**).

## Discussion

Genomic loci enriched with segmental duplications, including the IGK locus [14,23], lack complete and accurate information on population-level variation due to shortcomings of short-read sequencing [10,43,44].

Here, we use a SMRT long-read sequencing and analysis framework to characterize diploid IGK assemblies from 36 individuals of African, East Asian, South Asian, European, and Central American ancestry. These assemblies represent a significant advance, increasing the number of available curated assemblies 18-fold, from 4 [13–15,18] to 72 haplotypes. We identify a common SV ∼24.7 Kbp in length that includes a functional IGKV gene observed more frequently in AFR individuals relative to three other populations. Analysis of 47 functional IGKV genes revealed at least one novel allele for 33 of these genes, with 64 novel alleles identified in total. Fourteen of the IGKV genes showed evidence of inter-population allelic variation, including alleles unique to specific populations within this dataset. Finally, we demonstrate that a gene conversion event associates with a distinct IGK distal haplotype structure that is common in the population.

Databases of germline IG alleles are foundational for the analysis and interpretation of AIRR-seq data [44,45] and, more fundamentally, genotyping is required to determine the impact of IG germline variation on IG gene usage in antibody repertoires [6]. To our knowledge, the data presented here represent the first survey of human IGK haplotypes in an ancestrally diverse cohort. Novel IGKV allele sequences described in this study represent a ∼64% increase in IGKV alleles available in IMGT and have been made public in VDJbase and the Open Germline Receptor Database (OGRDB) as part of an ongoing effort to build a population-representative germline allele database [39,46,47] that can be leveraged for applications such as AIRR-seq analysis. The high prevalence of novel alleles in AFR individuals as compared to each other population reflects the ongoing need to generate genomic resources that are representative of the global population [42].

Deficits in existing resources for other forms of genetic variation were also observed in this dataset. For example, of the >3,000 common SNVs identified, ∼45% were not found in dbSNP. This is likely a reflection of the fact that locus complexity has hindered the use of high-throughput genotyping methods for cataloging IGK variants. We demonstrated that for most individuals in our cohort, over half of the SNVs called using our approach were missing from the 1KGP phase 3 genotype sets. This result alone implies that genetic disease association studies in the IGK locus have likely not effectively surveyed genetic diversity in this region.

Structural variation is common in the IG loci [4,6,7,14]; here, we identified a ∼24.7 Kbp insertion in the IGK distal region that included the functional gene IGKV1-NL1. Higher frequency of the insertion sequence in

AFR individuals is consistent with the observation that AFR populations harbor increased amounts of sequence in SD regions genome-wide [48,49]. Inclusion of this structural variant in future human genome references will be important for not only describing IGKV1-NL1 alleles, but also for genotyping all SNVs in this SV that can be tested for association with immune-related phenotypes. It is notable that the IGKV1-NL1 insertion was the sole SV identified in this cohort, indicating that SVs may be less characteristic of the IGK locus relative to IGH and IGL.

It is appreciated that SDs provide substrates for more frequent gene conversion, in some cases leading to enrichments of SNVs [43]. We previously reported the existence of two distinct haplotype structures within ∼16 Kbp of the IGK distal region; one of these haplotypes harbored a gene conversion with sequence at IGKV1D-13 exchanged for sequence at IGKV1-13 [14]. Here, we demonstrate that haplotypes harboring the gene conversion are common in the population and associate with increased SNV density (relative to the remainder of the IGK distal region) throughout ∼100 Kbp ranging from IGKV1D-17 to IGKV1D-43. The frequency of both the gene conversion haplotype and the IGKV1-NL1 SV were similar in EAS, SAS, and EUR populations but distinct among AFR individuals. Inter-individual haplotype variation in the IGK distal region was notable among AFR individuals, whereas most individuals from other populations were grouped in clusters (**Figure 4F**). These observations may reflect a high degree of genetic diversity in AFR populations that is reduced by forms of genetic drift, such as bottleneck effects, that occurred in non-AFR populations [50,51].

Alternatively, the haplotype structures observed in AFR individuals may reflect selection of alleles that elicit expression of light chain antibody repertoires which confer an advantage when confronted with pathogen-mediated diseases. Given the cohort analyzed here was relatively small, analysis of additional haplotypes will be needed to determine the extent of inter-population variation with respect to haplotype structure in IGK proximal and distal regions.

Considering that IGH germline variants are associated with usage of IGH genes in expressed antibody repertoires [6], we hypothesize that the usage of IGK genes will also likely be associated with SNVs and SVs identified here. AIRR-seq studies focused on IGL and IGK have shown inter-individual variation in germline gene usage and alternative splicing in expressed light chain repertoires [52]. The data and analytical framework presented here lay a foundation for testing the impact of IGK locus-wide (coding and non-coding) germline variants on variation in the expressed antibody repertoire in diverse human populations. This will expand our understanding of the collective roles of IG germline variation across the IGH, IGL and IGK loci in antibody diversity and function, including potential genetic interactions between these complex gene regions and their co-evolution [21,23]. This will ultimately be critical for identifying coding alleles and haplotype variation as determinants of disease susceptibility and clinical phenotypes, including autoimmunity, cancer, infection, and vaccine efficacy [9,11,21,24,53–65].

### Data Availability

Novel alleles are provided in Table S3. Mapped sequence data and assemblies will be made available at https://vdjbase.org/ with sample metadata. Raw data are undergoing submission to BioProject PRJNA555323.

## Acknowledgements

We acknowledge and thank William Gibson for productive insights regarding IG light chain genetic variation, and we also acknowledge Zachary VanWinkle for assistance with manuscript preparation.

## Supplementary Figure Legends

**Supplementary Figure 1.**
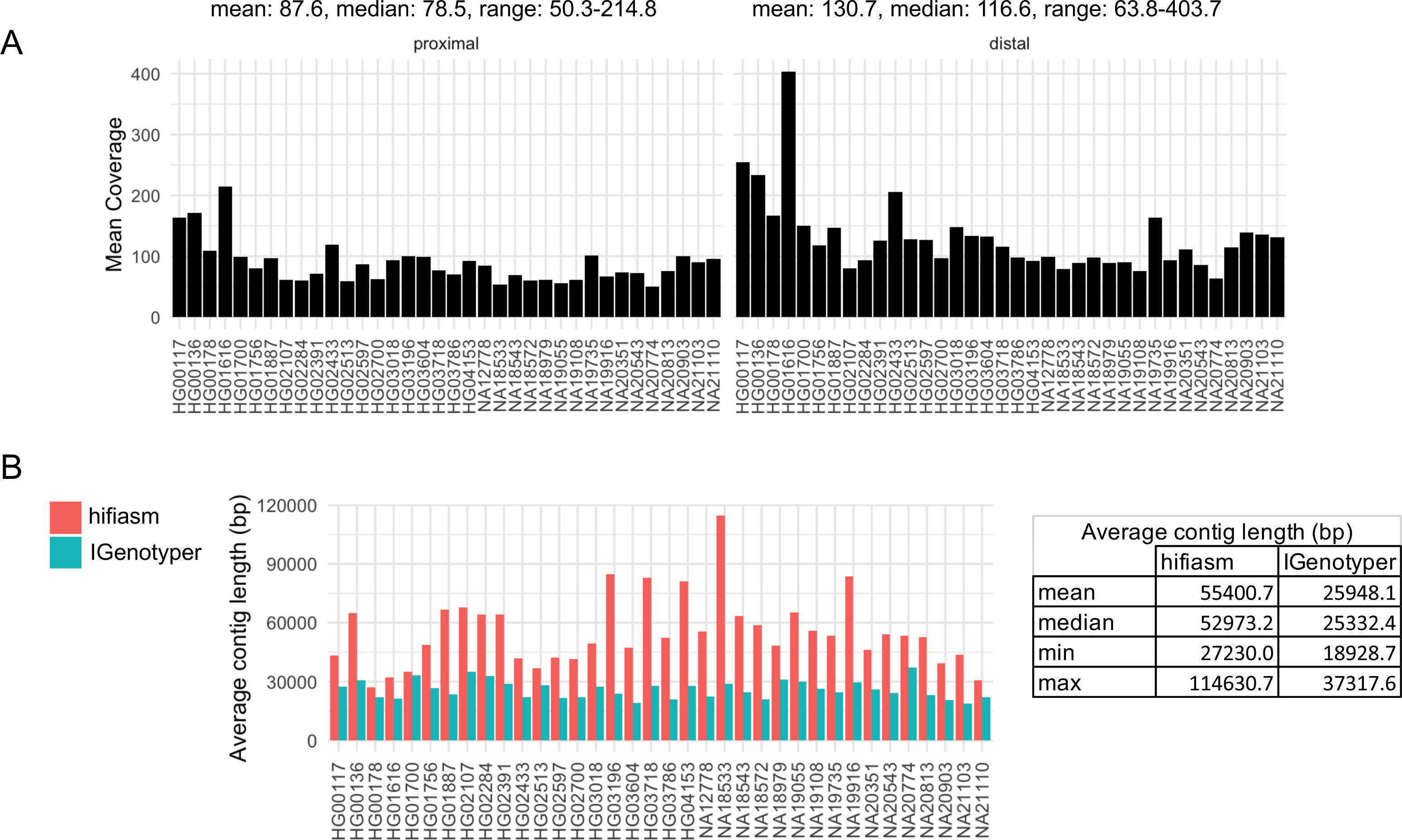
**(A)** HiFi read coverage across the IGK locus for all samples in the cohort. **(B)** Average length of contigs generated by hifiasm and IGenotyper for each sample.

**Supplementary Figure 2.**
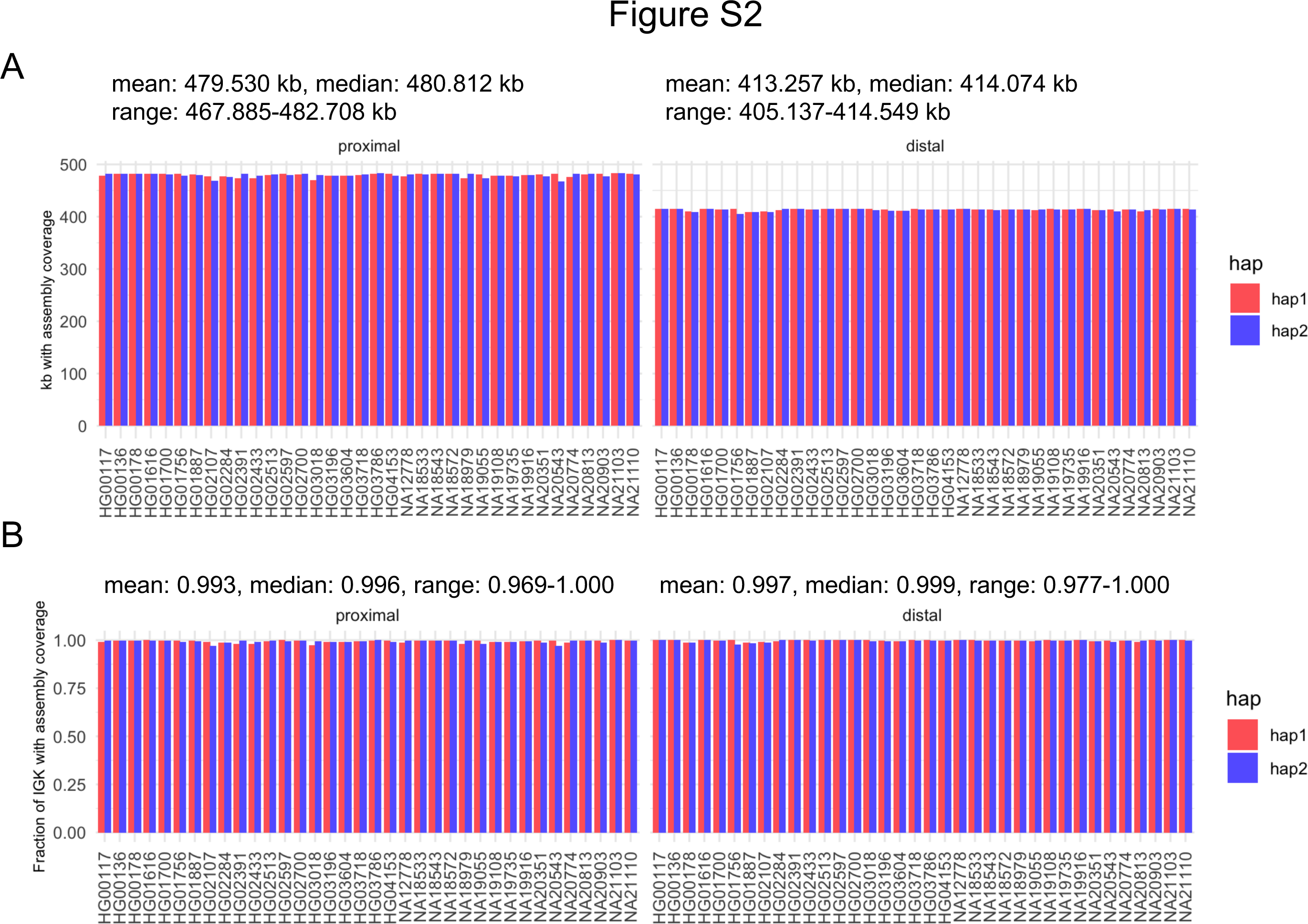
**(A)** Number of reference bases with curated assembly coverage. **(B)** Fraction of IGK proximal and distal regions with curated assembly coverage.

**Supplementary Figure 3.**
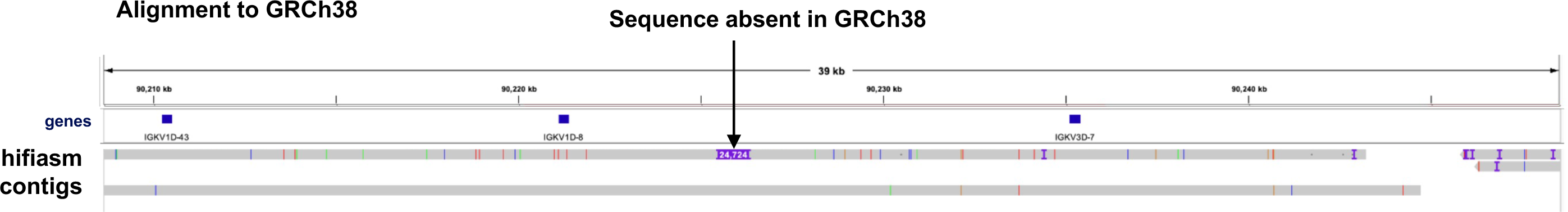
IGV screenshot of hifiasm-generated contigs from an AFR sample mapped to the GRCh38 genome assembly. One of the contigs (one haplotype) has a ∼24.7 Kbp insertion between IGKV1D-8 and IGKV3D-7.

**Supplementary Figure 4.**
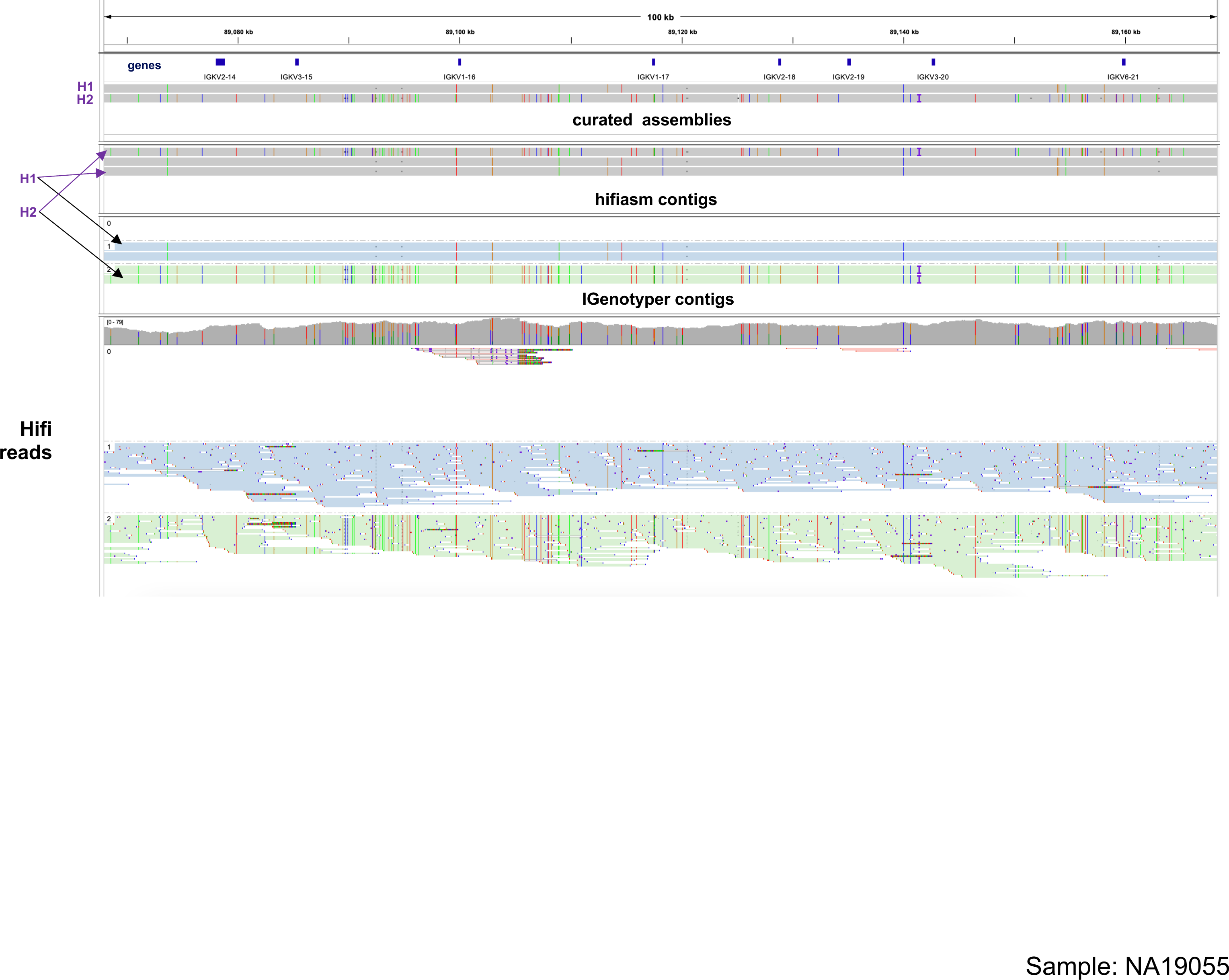
IGV screenshot of 100 Kbp in the IGK proximal region with curated assemblies, phased HiFi reads as well as phased contigs generated by IGenotyper and hifiasm aligned to our custom reference. Purple arrows indicate haplotype (“H1” and “H2”) contigs selected for curated assembly.

**Supplementary Figure 5.**
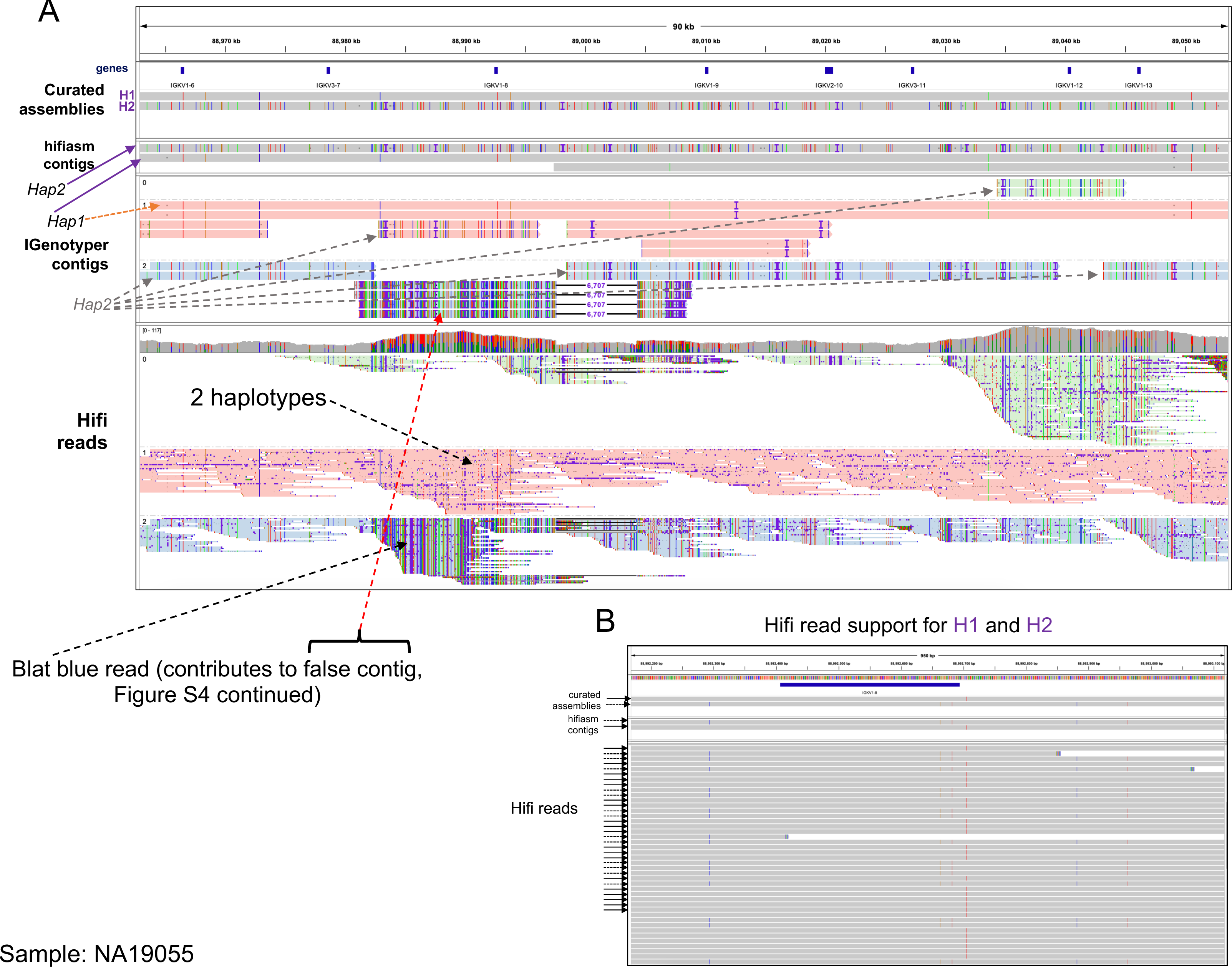
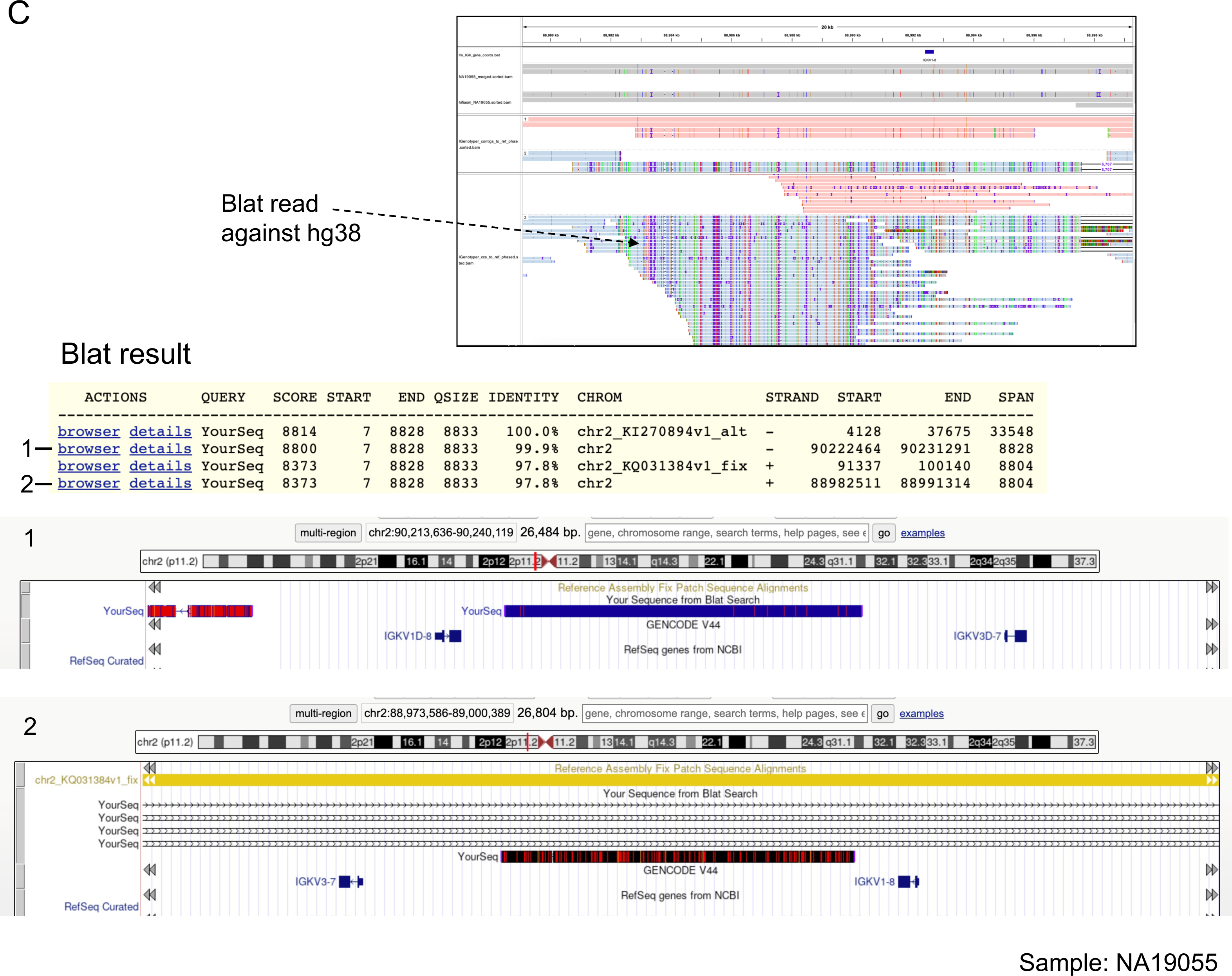
(A) IGV screenshot showing curated assemblies (H1 and H2), hifiasm contigs, IGenotyper contigs, and HiFi reads aligned to our custom reference. Purple arrows indicate contigs selected for curated assemblies. An IGenotyper contig (orange arrow) congruent with the “Hap1” contig generated by hifiasm (purple arrow). The “Hap2” contig generated by hifiasm (purple arrow) is identified as discontiguous contigs generated by IGenotyper (gray arrows). The red arrow indicates a group of assemblies that likely result from mismapped reads. The black arrow indicates a read that was queried against the GRCh38 genome (continued). **(B)** IGV screenshot of curated assemblies, hifiasm contigs, and HiFi reads, zoomed in to a region containing IGKV1-8 in panel (**A**). Two SNP patterns in HiFi reads (dashed and solid arrows) are consistent with hifiasm contigs and curated assemblies. **(C)** IGV screenshot showing the aforementioned read, which was queried against the GRCh38 genome using BLAT (UCSC Genome Browser). The best match was to the IGK distal region, between IGKV1D-8 and IGKV1D-7, and the second-best match was to the IGK proximal region, between IGKV3-7 and IGKV1-8.

**Supplementary Figure 6.**
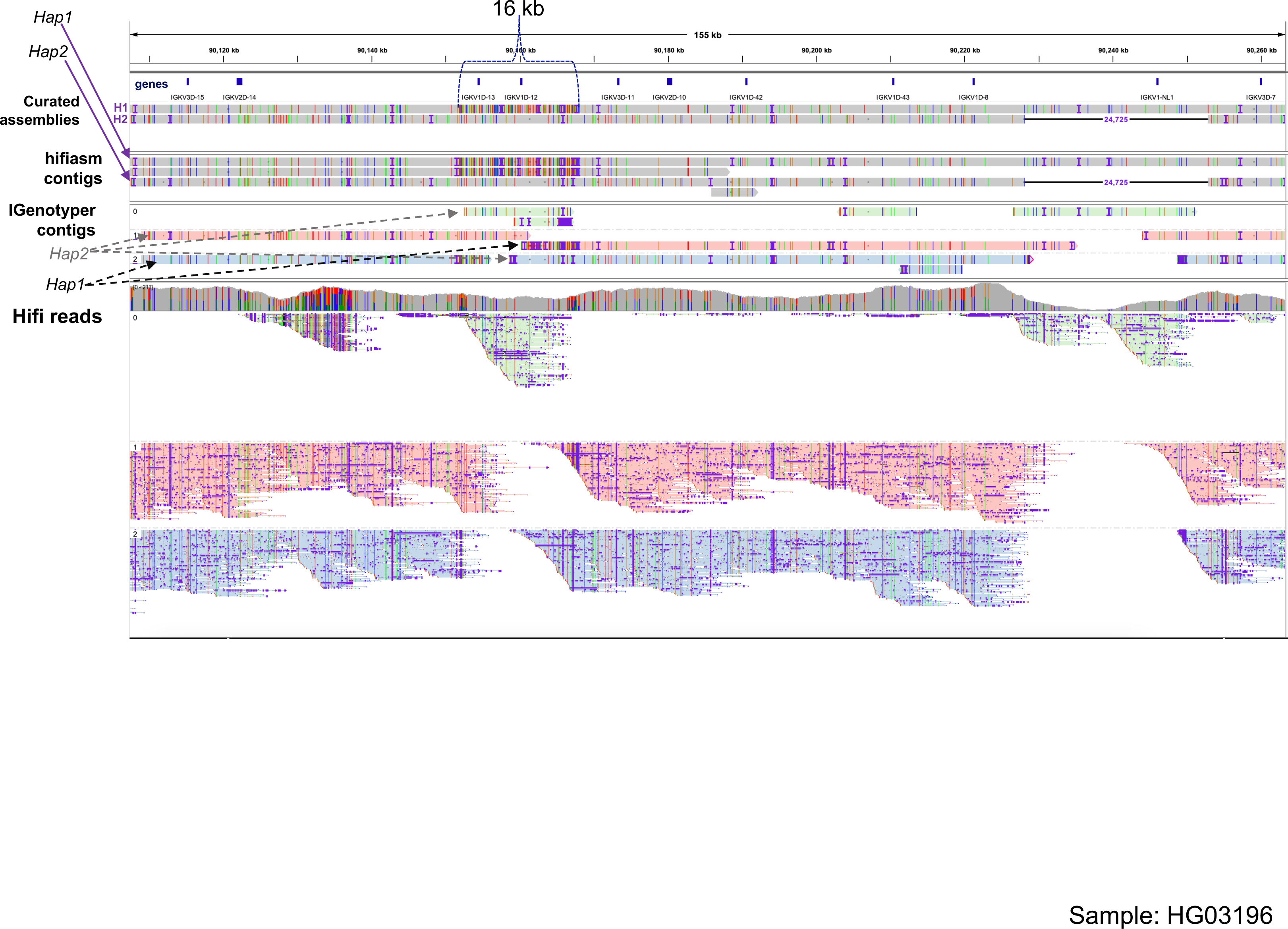
IGV screenshot of the IGK distal region spanning IGKV3D-15 to IGKV3D-7, which includes the IGKV1- 13/IGKV1D-13; IGKV1-12/IGKV1D-12 gene conversion (∼16 Kbp region) and structural variant that includes the gene IGKV1-NL1. Shown are curated assemblies (H1 and H2), hifiasm contigs, IGenotyper contigs, and HiFi reads aligned to our custom GRCh38 reference. Purple arrows indicate hifiasm contigs selected for curated assemblies; “Hap1” (H1) includes the gene conversion event. IGenotyper-generated contigs corresponding to “Hap1” and “Hap2” are fragmented across multiple contigs; hifiasm generated uninterrupted contigs for “Hap1” and “Hap2’’ in this region.

**Supplementary Figure 7.**
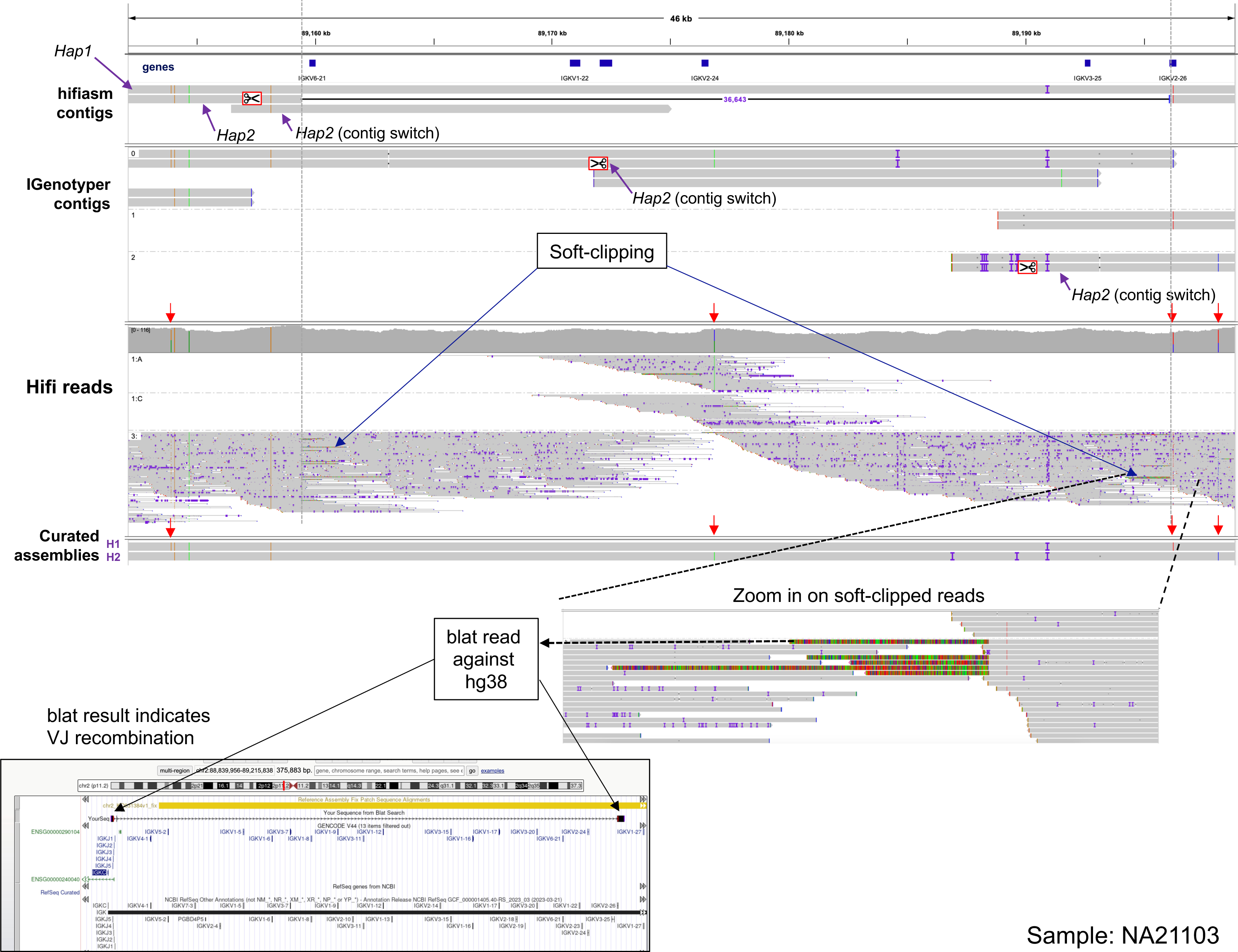
IGV screenshot of 46 Kbp in the IGK proximal region with hifiasm-generated contigs, IGenotyper contigs, HiFi reads, and curated assemblies aligned to our custom reference. A minority of HiFi reads have soft-clipped bases that correspond to the start and end of a deletion present in a hifiasm contig. The indicated read with soft-clipped bases (dashed arrow) was queried against the GRCh38 genome using BLAT; the best match resulted in a split alignment, with part of the read matching to a V gene (IGKV2-26) and the remainder of the read mapping to the IGKJ gene region; this is a signature of V(D)J recombination in LCLs [7,8], with the presence of reads without soft-clipping explained by cell line polyclonality [66].

**Supplementary Figure 8.**
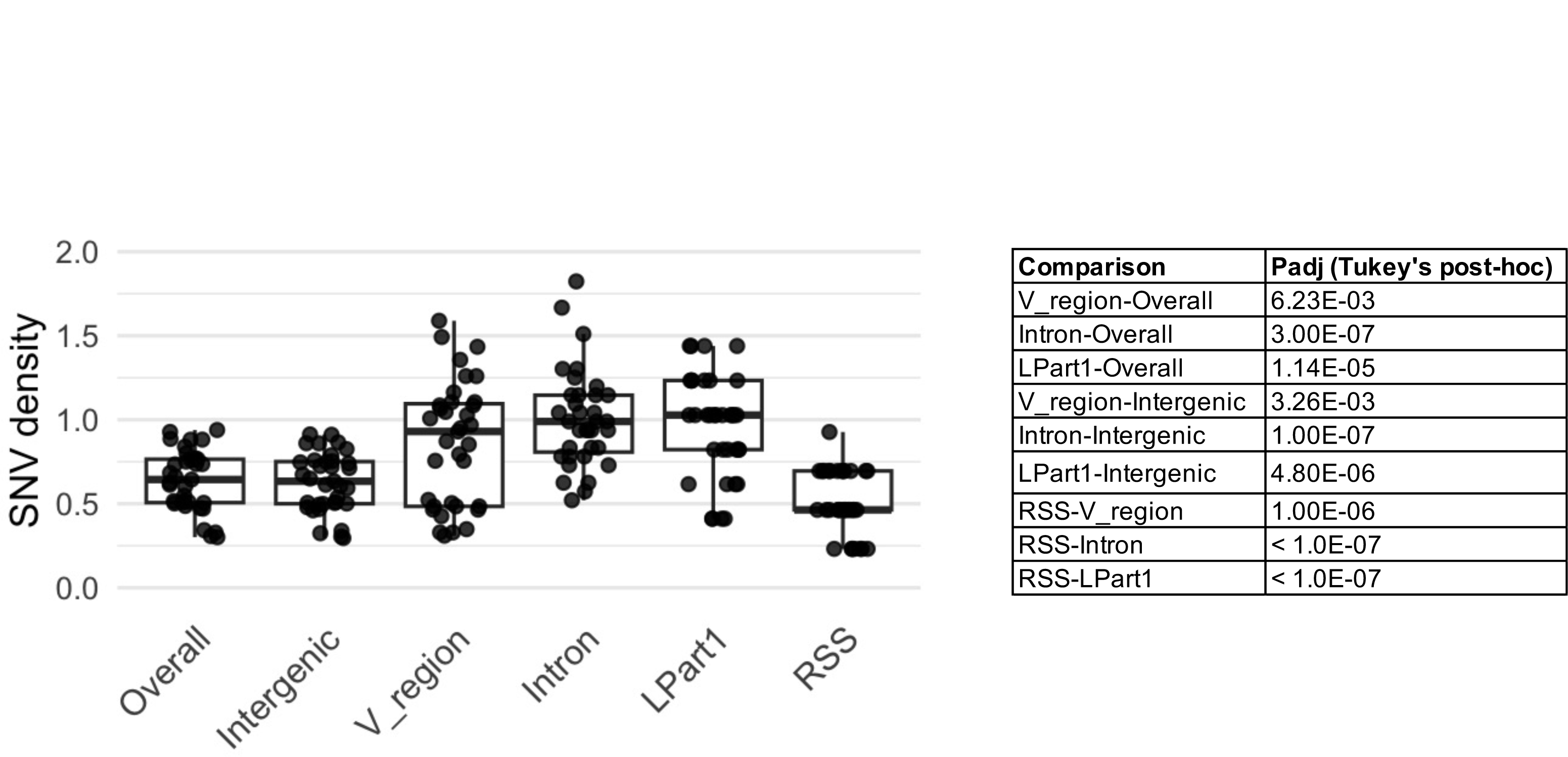
SNV densities for indicated genomic features for 35 samples (data points). Table indicates adjusted P-values from Tukey’s post-hoc test. Comparisons not shown have adjusted P-values > 0.05.

**Supplementary Figure 9.**
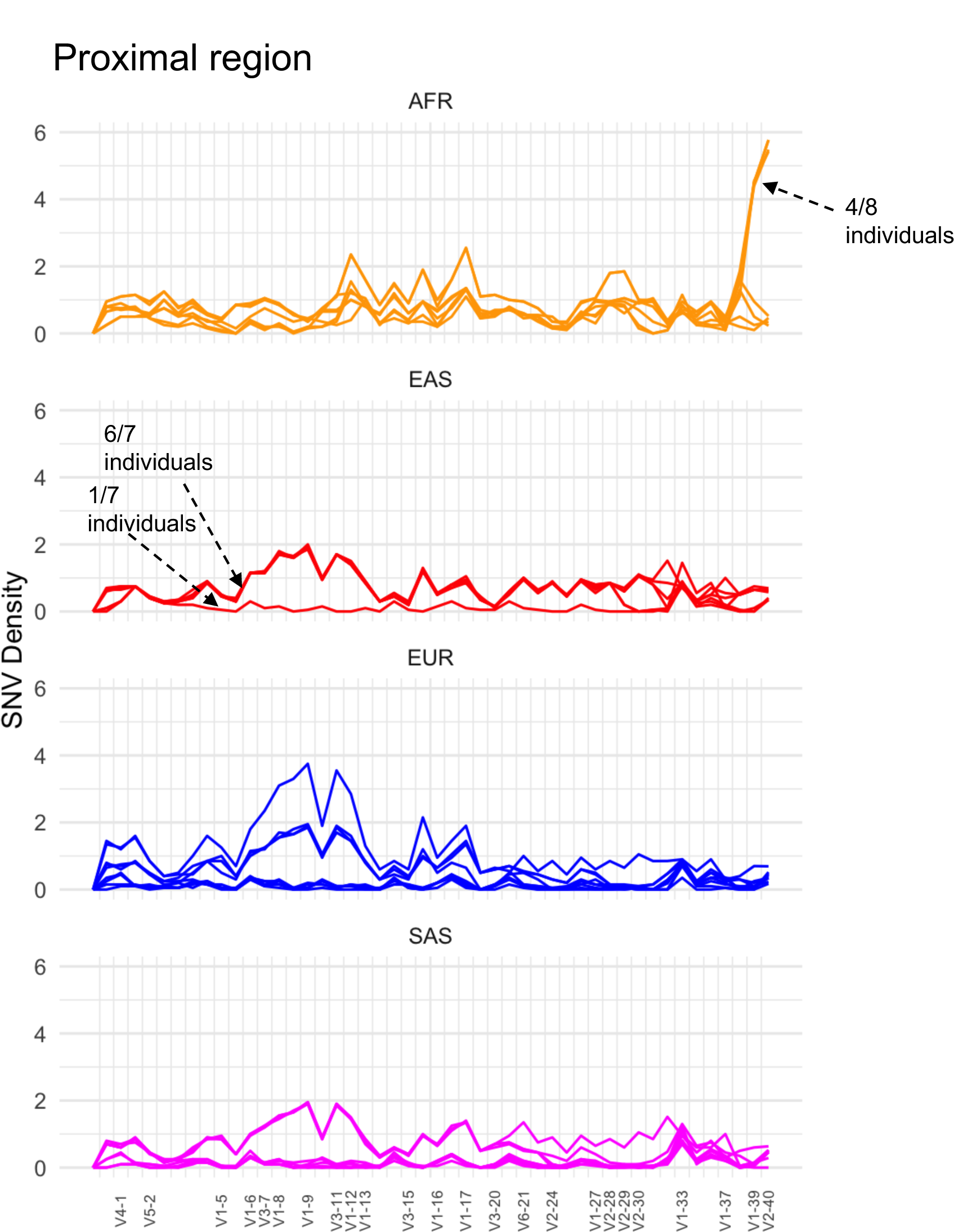
SNV densities in 10 Kbp windows along the IGK proximal region. Each line represents an individual. Individuals are grouped according to population. Samples are indicated in selected regions where lines (samples) are overlapping. IGKV gene positions are indicated.

**Supplementary Figure 10.**
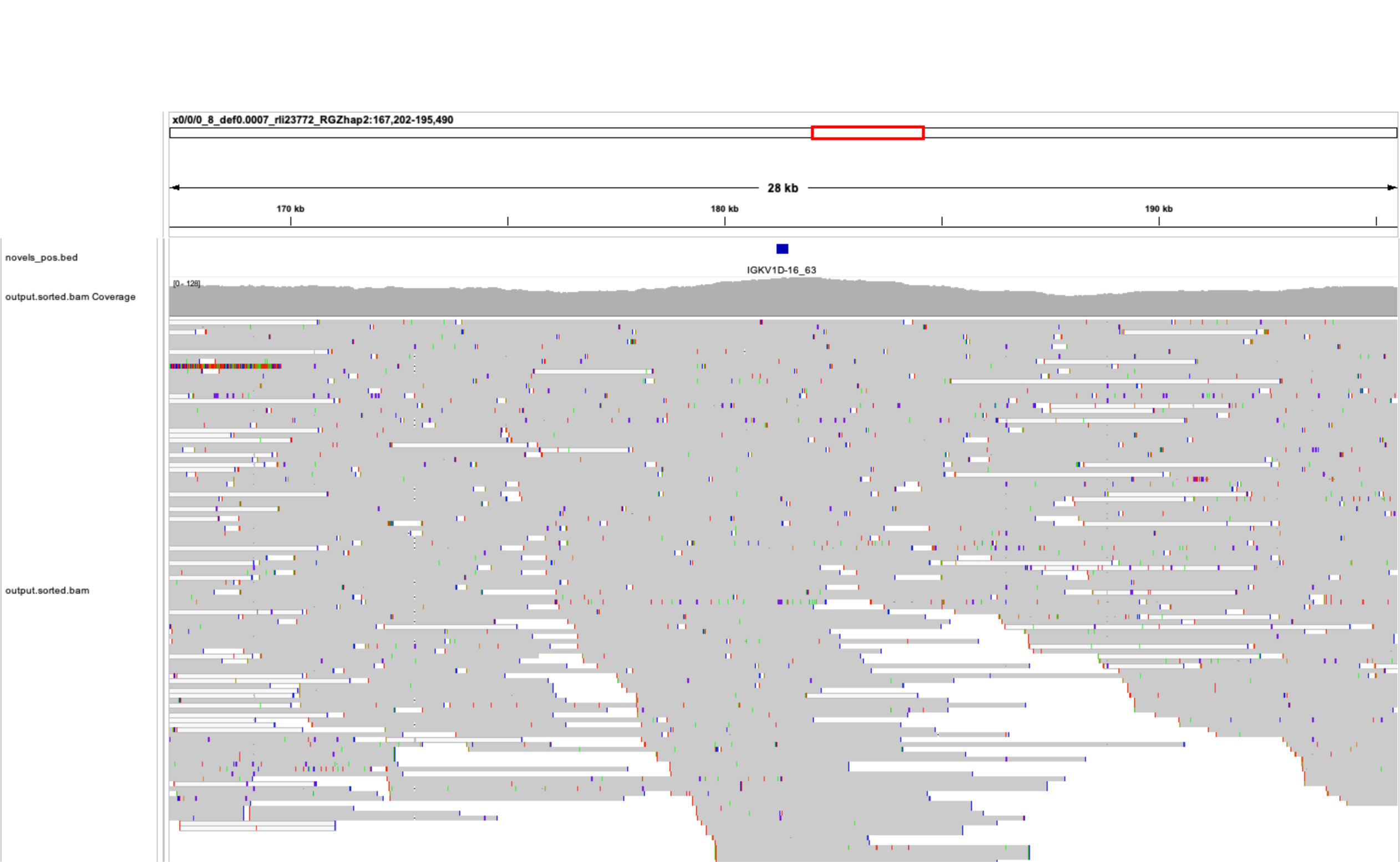
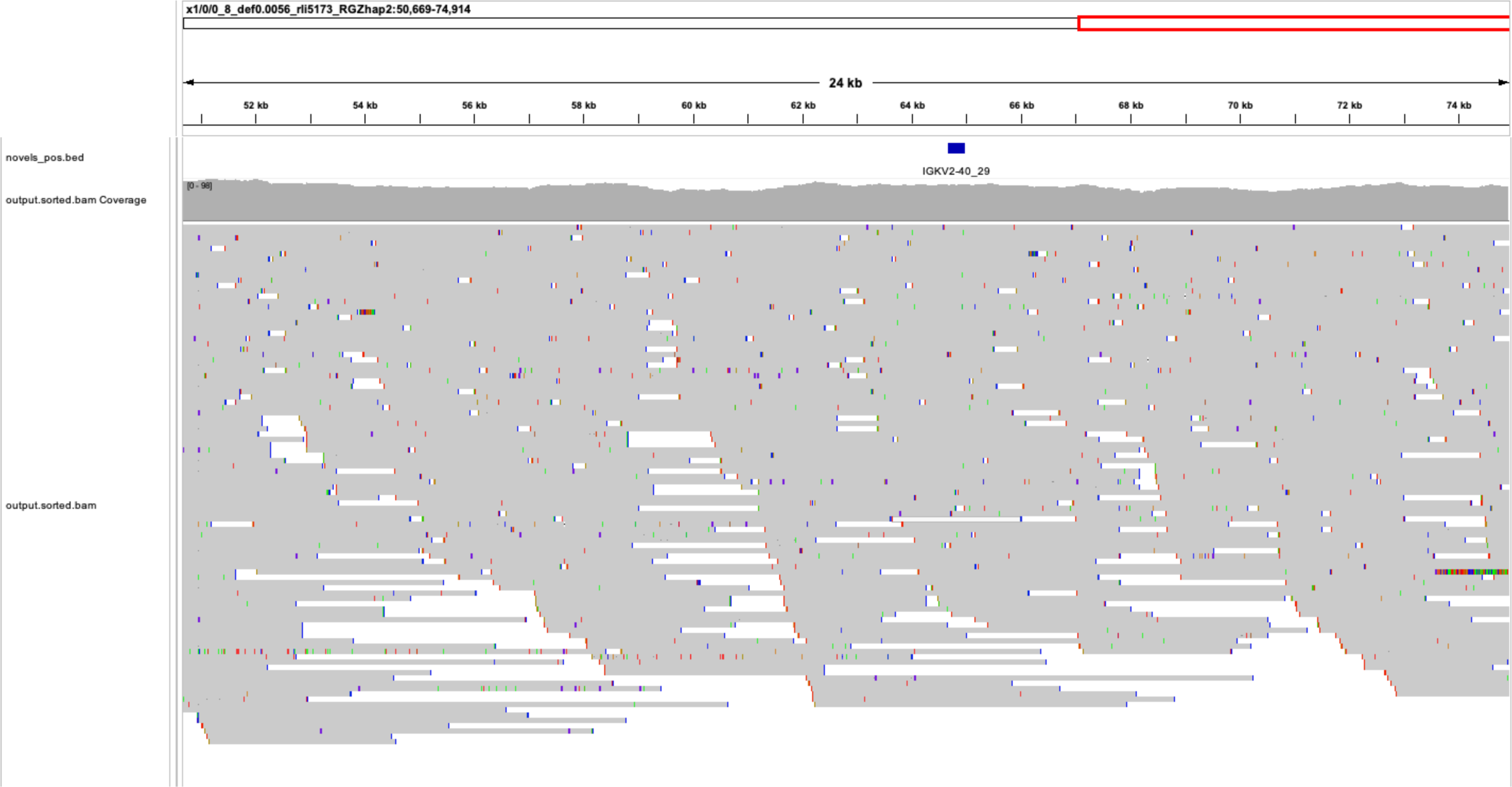
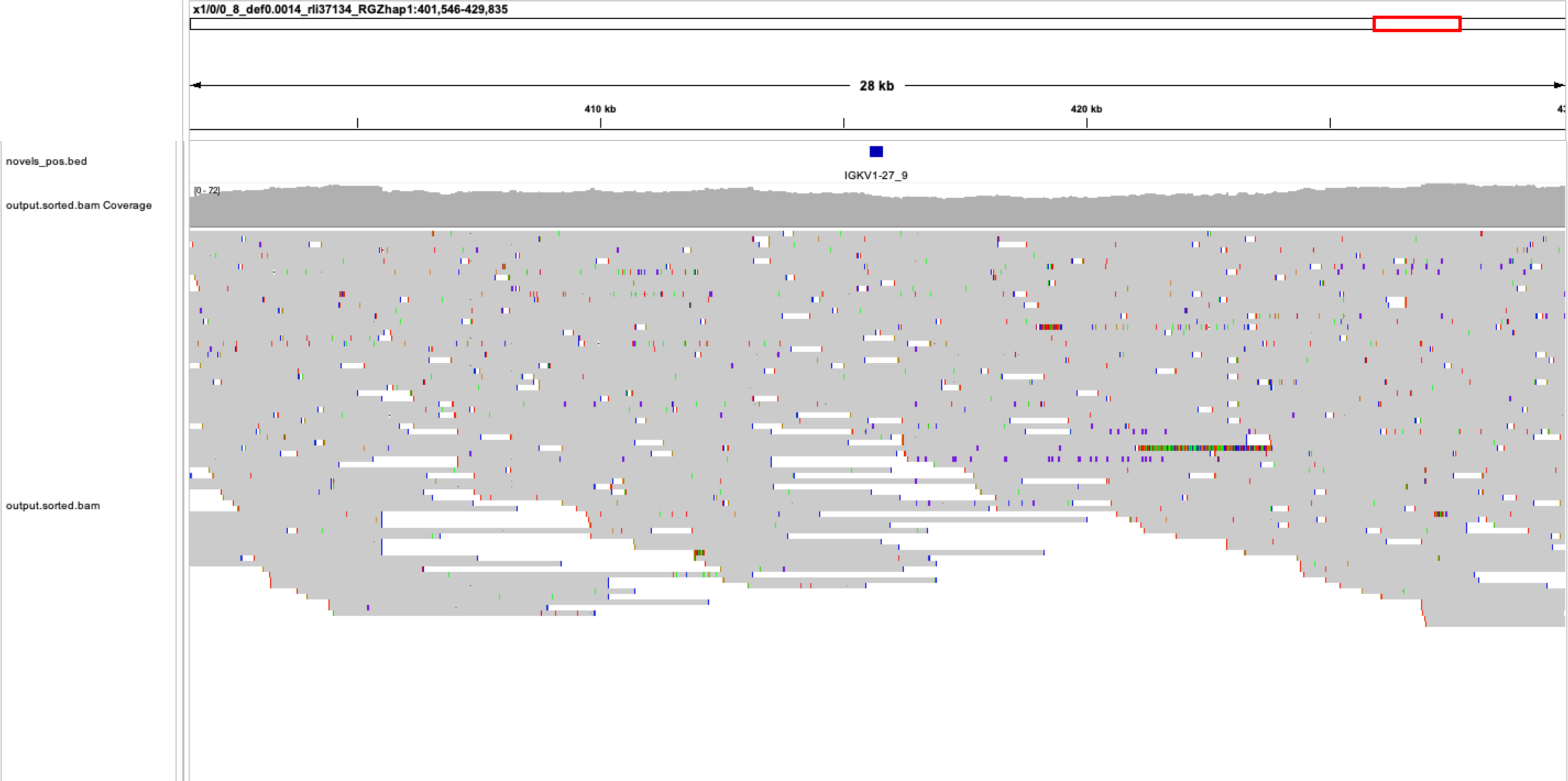
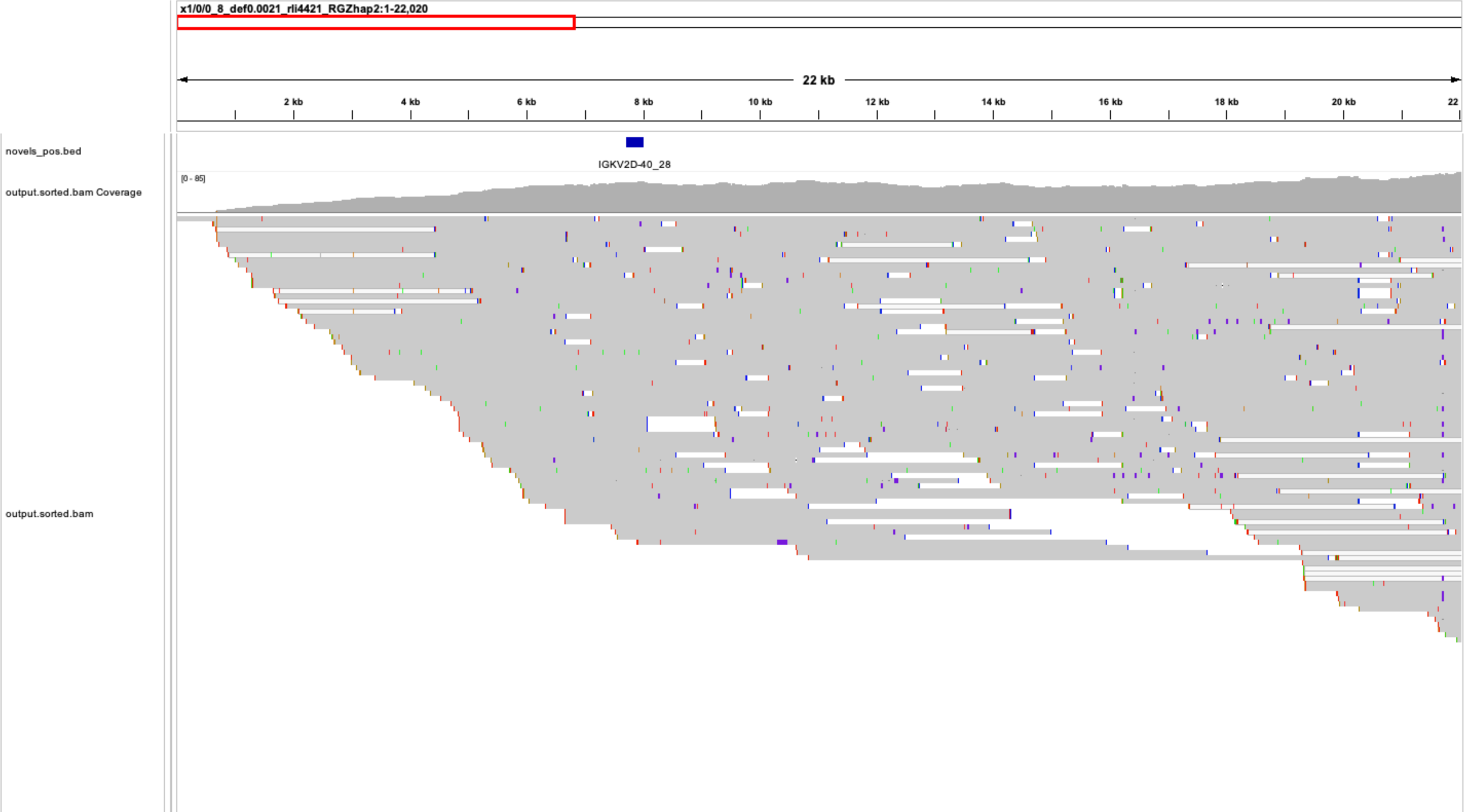
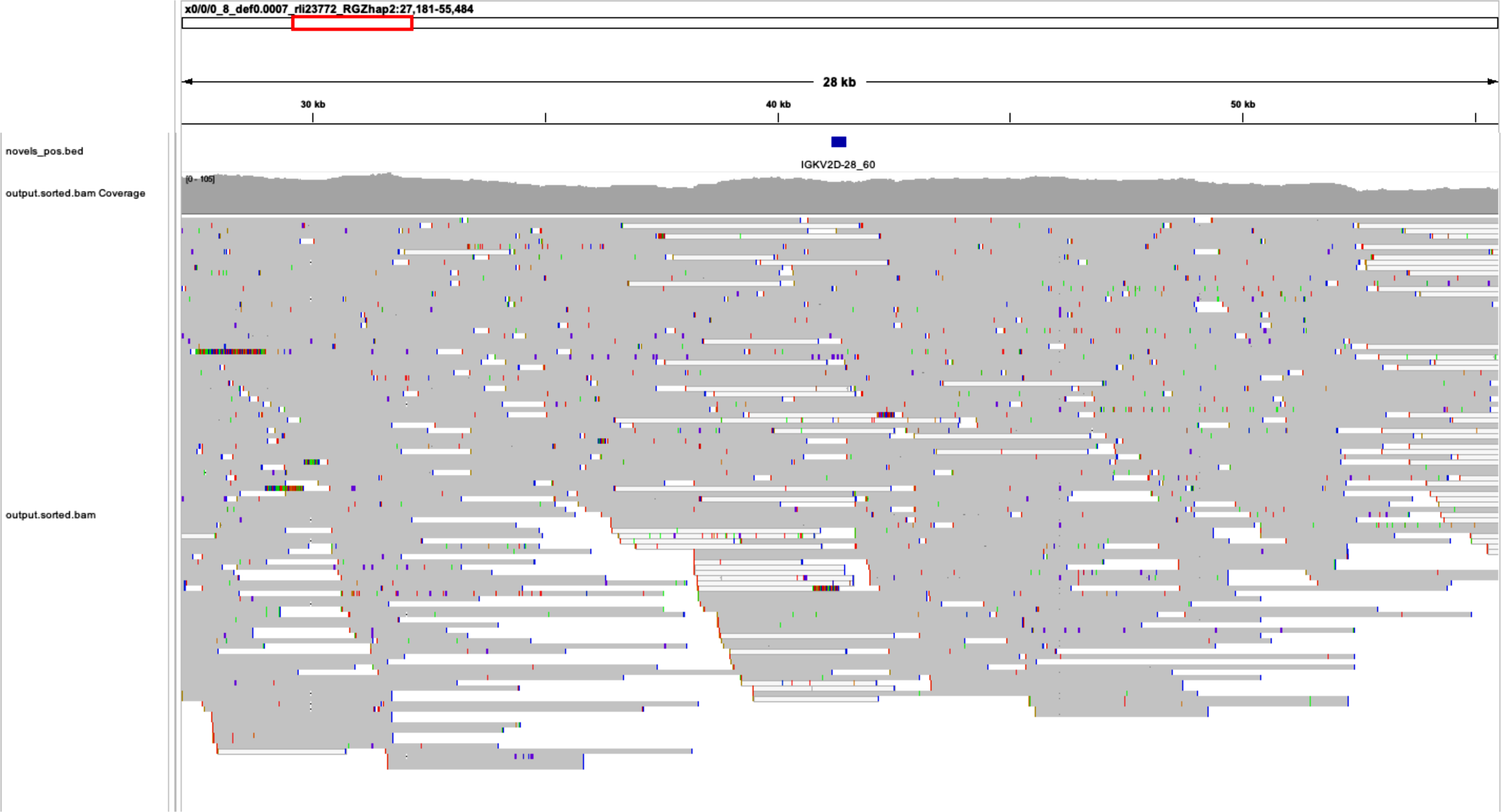
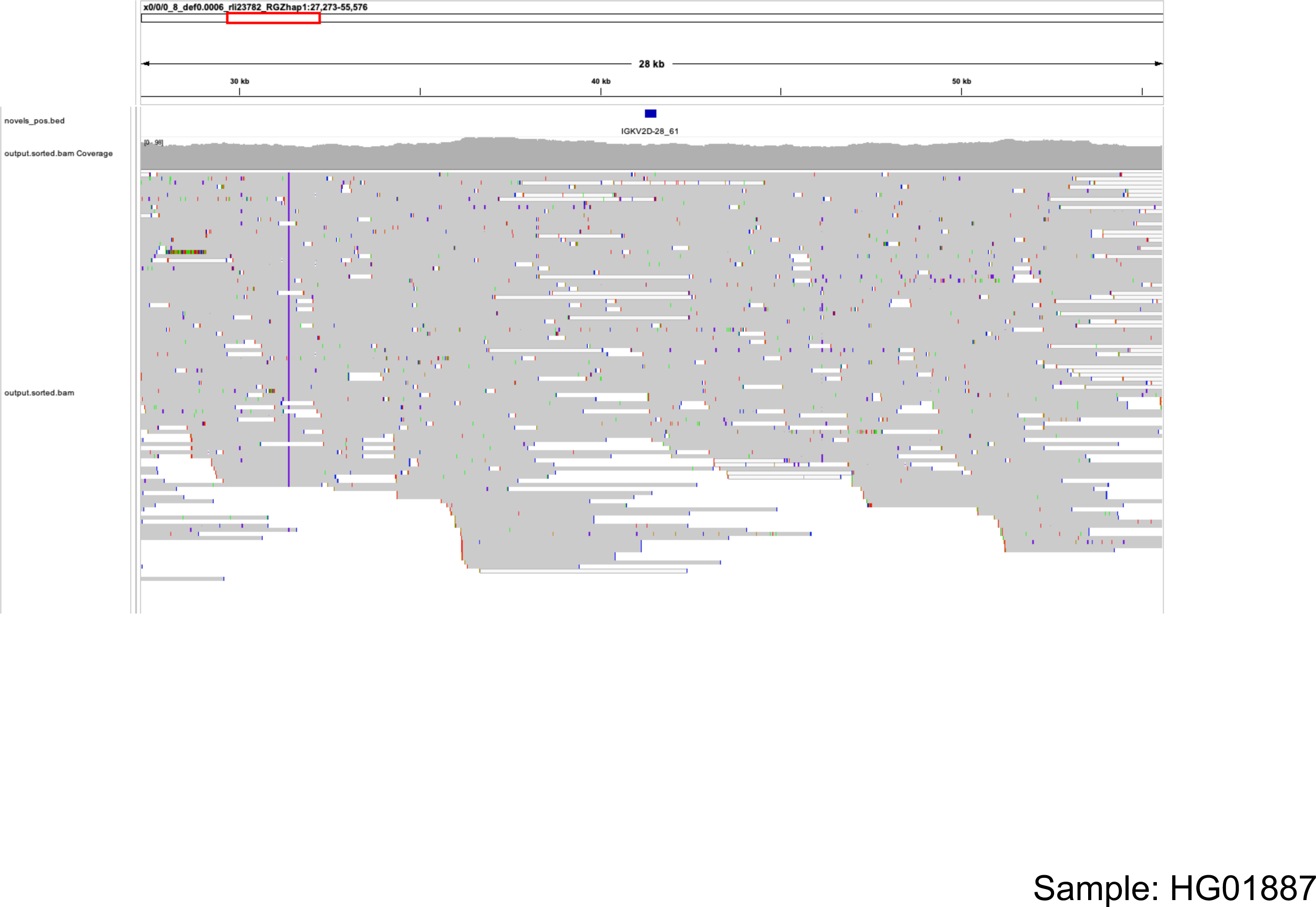
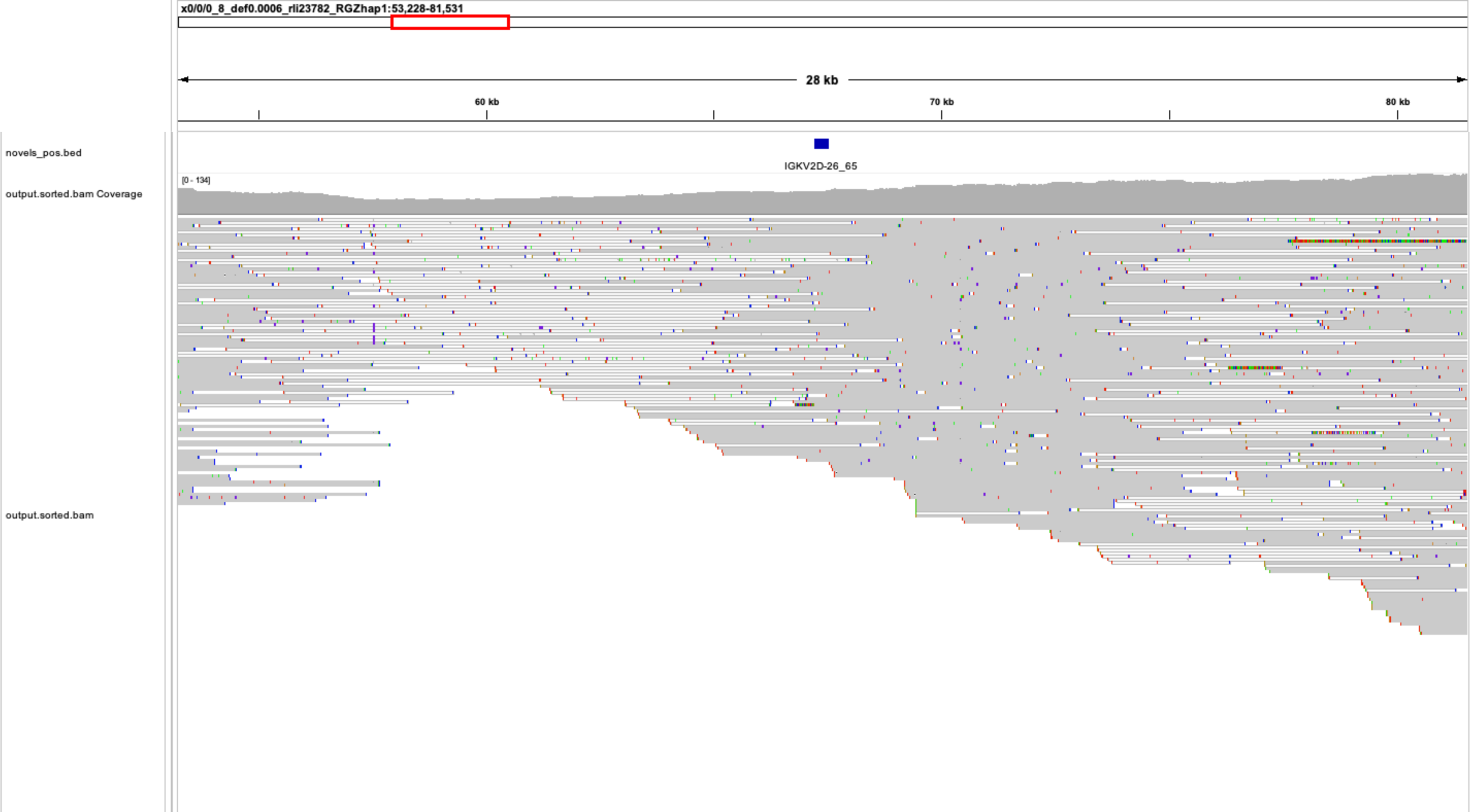
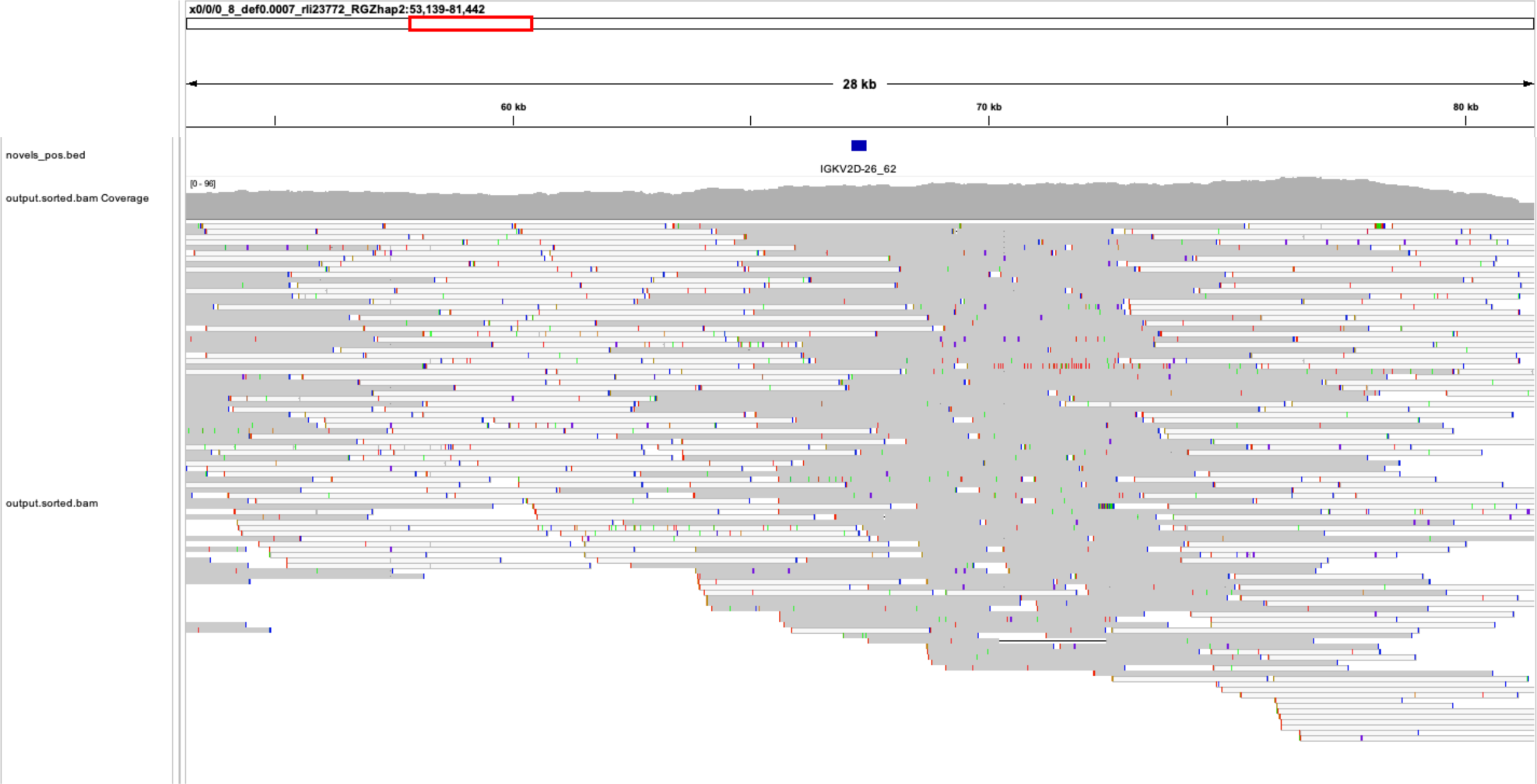
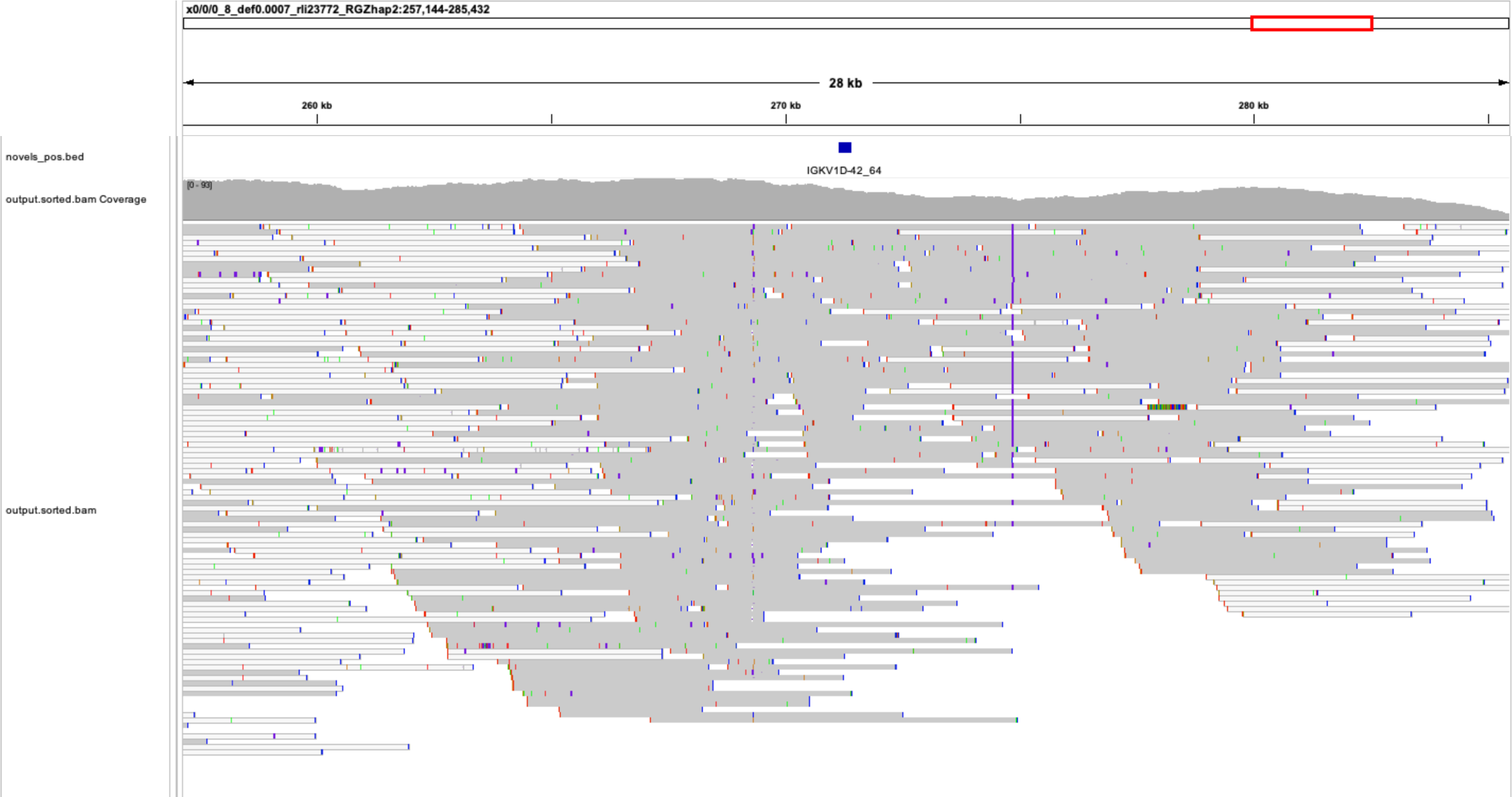
IGV screenshots of novel alleles from the sample HG01887 aligned to the individual’s IGK-personalized reference (see **Materials and Methods**). Also shown are HiFi reads aligned to the individual’s IGK- personalized reference. HiFi reads support (i.e. do not show mismatches with) the contigs selected for diploid IGK assembly.

